# Wild type bone marrow-derived microglia-like cells mitigate strain-dependent neuropathology in Csf1r mutant mice

**DOI:** 10.1101/2025.01.16.633478

**Authors:** Isis Taylor, Omkar L. Patkar, Yajun Liu, Sebastien Jacquelin, Emma K. Green, Ginell Ranpura, Dylan Carter-Cusack, Nikita Rosendahl, Adam Ewing, Nyoman D. Kurniawan, Mosi Li, Deepali Vasoya, Xin He, Owen R. Dando, Peter Kind, Giles E. Hardingham, Barry M. Bradford, Neil A. Mabbott, Lucas LeFevre, Clare Pridans, Kim M. Summers, Katharine M. Irvine, David A. Hume

## Abstract

Germ-line deletion of a conserved enhancer (FIRE) in the mouse *Csf1r* locus (*Fireko*) causes congenital absence of microglia. The current study analyses the effect of microglial deficiency on gene expression during postnatal brain development and defines the effect of mouse genetic background on the *Fireko* phenotype and age-dependent pathology. In total RNA-seq data from multiple brain regions of juvenile wild-type and *Fireko* mice, transcripts associated with endosome/lysosome function and other transcripts strongly enriched in isolated microglia were not affected indicating compensatory expression by other cell types. The *Fireko* mutation on a C57BL/6J background led to perinatal lethality and hydrocephalus (HC) in surviving pups that was overcome in F2 intercross with CBA or BALB/c. The absence of microglia in *Fireko* mice did not affect motor activity, behaviour, or myelination but aged mice developed astrocytosis and subsequent severe calcification in the thalamus. Intraperitoneal transfer of WT bone marrow cells (BMT) repopulated the brain with microglia-like cells that expressed damage-associated transcripts and proteins. BMT prevented calcification in the thalamus but neither increased nor decreased markers of astrocyte or oligodendrocyte activation. The results suggest that the damage-associated microglial phenotype is a protective response to injury rather than a cause of neuropathology.

**eTOC Summary:** In this study Taylor et al. analyse a mouse model of a human genetic disease that leads to neurodegeneration. They show that the loss of key cells in the brain (microglia) can be reversed by transfer of donor marrow cells without toxic conditioning leading to reduced brain pathology.

## Introduction

Early in vertebrate development, the brain is populated by phagocytes derived from the yolk sac and fetal liver. These give rise to microglia and brain-associated or border macrophage (BAM) populations (Prinz et al., 2021; Van Hove et al., 2019). In the postnatal period, microglia and a further wave of recruited monocytes (Askew et al., 2017; Carrier et al., 2024; Perry et al., 1985) are involved in phagocytic clearance of excess neurons and synapses. Together, these populations are proposed to support or modify neurogenesis, oligodendrocyte function and myelination, and astrocyte function (Carrier et al., 2024; Chen et al., 2020; Leong and Ling, 1992; Mehl et al., 2022; Paolicelli et al., 2022; Prinz et al., 2021; You et al., 2024).

The differentiation, proliferation and survival of tissue macrophage populations including microglia depends upon signals from the CSF1R tyrosine kinase receptor (Hume et al., 2020). The *Csf1r* locus contains an intronic super-enhancer (Fms intronic regulatory element; FIRE) which is conserved from reptiles to humans (Hume et al., 2017). Mice with a germline deletion of this element are completely microglia-deficient, and partially deficient in BAM (Rojo et al., 2019). They are almost entirely macrophage deficient in the embryo (Munro et al., 2020) and lack microglia and specific tissue macrophage populations in the adult (Liu et al., 2026; Rojo et al., 2019).

Heterozygous dominant and homozygous recessive mutations in the human *CSF1R* gene are associated with neuropathology (Dulski et al., 2024; Guo and Ikegawa, 2021). The disease spectrum, now referred to as CSF1R-related leukoencephalopathy (CRL) or disorder (CRD) is associated with profound microglial deficiency and dysregulation of astrocyte and oligodendrocyte function (Du et al., 2025). *Fireko* mice have been analysed as a model for the effect of microglial deficiency seen in patients. McNamara *et al*. (McNamara et al., 2023) reported age-dependent impacts on oligodendrocyte function and myelin integrity in *Fireko* mice and two recent studies described calcification and neuronal loss in the thalamus that could be prevented by intracerebral microglial transplantation (Chadarevian et al., 2024; McNamara et al., 2023).

In inbred *Fireko* mice, intraperitoneal injection of wild-type bone marrow cells is sufficient to fully repopulate vacant territories, including the brain, without contributing to the blood monocyte pool (Liu et al., 2026). Here we examine the impact of long term chimerism on the development of neuropathology including calcification of the thalamus. Notwithstanding the expression of disease-associated microglial genes in donor-derived cells, our results indicate that bone marrow-derived microglia can have a therapeutic benefit in treatment of microglia deficiency.

## Results

### Selective loss of microglial-specific transcripts throughout the brain in Fireko mice

Deconvolution of total RNA-seq data provides an alternative approach to quantitation of cell-type and process-specific transcriptomic signatures in complex tissues without the requirement for cell isolation (Huuki-Myers et al., 2025). To identify possible mechanisms of compensation for the absence of microglia or early indicators of pathology we analysed bulk RNA sequencing on cortex, hippocampus, and striatum from two-week-old mice, and cortex, hippocampus, striatum, cerebellum, and olfactory bulb from six-week-old *Fireko* mice and littermate controls on the original C57BL/6J x CBA (3:1) mixed background (*Fireko*^Edin^). The full datasets are provided in **Table S1** and **S2**, respectively, with conventional differential expression analysis for each region. As in our previous analysis in *Csf1r^-/-^*rats (Patkar et al., 2021) the RNA-seq data were analysed using the network analysis tool *BioLayout* and the clusters of transcripts with correlated expression and their average expression profiles are provided (**Table S3**).

As expected, microglia-specific transcripts were greatly reduced in the brains of *Fireko*^Edin^ mice. **Cluster 11** contained around 250 transcripts, including *Csf1r*, with the shared pattern of almost complete depletion in every brain region. **Cluster 30** showed a similar pattern, except that these transcripts were affected to a lesser extent by the *Csf1r* mutation (**Figure 1A**). The relative abundance of microglia inferred from the average expression of **Cluster 11** differed between brain regions, being lowest in cerebellum, but the set of co-expressed transcripts showed no evidence of the brain region specificity reported by others (Tan et al., 2020).

**Figure 1.**
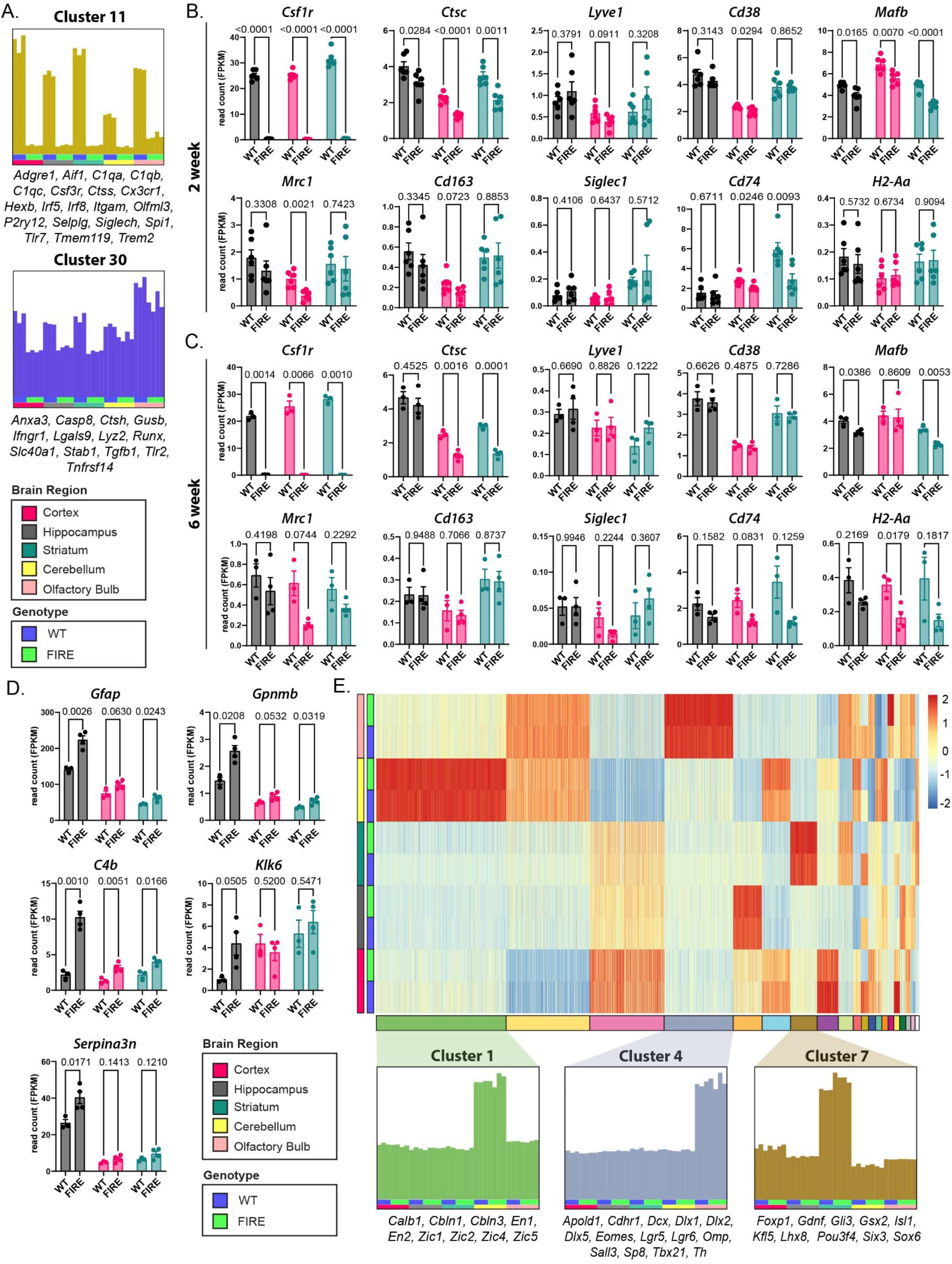
Transcriptomic analysis of the effect of the *Fireko* on juvenile mouse brain. RNA-seq data were generated from three brain regions of two-week-old (n=6/group) and five brain regions from six-week-old (n=3/4 per group) WT and *Fireko* mice on the C57BL/J x CBA genetic background (*Fireko*^Edin^). (A) Data obtained from six-week-old mice were clustered using the network analysis software *BioLayout* (full dataset provided in Table S3). Histograms show the average profile of the two large clusters correlated with *Csf1r* genotype. (B-C) RNA expression of select brain-associated macrophage markers in two-week-old (B) and six-week-old (C) hippocampus, cortex and striatum. (D) RNA expression of select markers of astrocyte or oligodendrocyte dysregulation in the same regions of six-week-old mice. Graphs show mean ± SEM, and each point is an individual animal. Differences between genotype were assessed by individual unpaired t tests with Welch’s correction. P values are reported on each graph. (E) Conventional heat map sorted by tissue and genotype (y axis) and cluster number (x axis) as indicated. The average expression profiles of three brain region-specific clusters identified by Biolayout, and representative marker genes, are shown below.

Adult *Fireko* mice retain brain-associated macrophages (BAM) (Rojo et al., 2019). In the analysis of *Fireko*^Edin^ pups at both two weeks and six weeks of age, transcripts encoding BAM markers (Van Hove et al., 2019) (*Ctsc, Lyve1, Cd38, Mafb, Mrc1, CD163, Siglec1, Cd74, H2-Aa*) remained detectable in all brain regions (**Table S1**; **Table S2**). The effect of the *Fireko* on individual markers was variable; *Lyve1, Cd38, Cd163* and *Siglec1* were unchanged where the other markers were marginally reduced (**Figure 1B, C).** Increased detection of *Gfap* and *Gpnmb* in *Fireko* cortex, hippocampus, and striatum at six weeks of age (**Figure 1D**; **Table S2**) provides an early indication of increased astrocyte reactivity. Previous single cell RNA-seq analysis of juvenile *Fireko* brains identified increased expression of *Serpina3n*, *C4b* and *Klk6* in a subset of oligodendrocytes (Munro et al., 2024). Of these markers, *C4b* was the most consistently upregulated in all brain regions in our bulk RNA-seq data (**Figure 1D**).

If the absence of microglia altered the relative abundance or expression signature of any cell type or biological process, we would detect a genotype-specific co-expression cluster. The microglia-related clusters provide a positive control and notably there are no transcripts within **Cluster 11** that are not highly-expressed by isolated microglia (Summers et al., 2020). **Figure 1E** shows expression values for genes present in the top 20 clusters and clearly delineates co-expression patterns present in each cluster. Aside from **Clusters 11** and **30**, other clusters were brain region-specific, and none was affected by the *Fireko* mutation **(Figure 1E**). For example, **Cluster 1**, strongly enriched in the cerebellum relative to all other brain regions, contains genes encoding known transcriptional regulators of cerebellar differentiation (e.g. *En1, En2; Zic1, Zic2, Zic4, Zic5)* as well as markers of cerebellar cell types (e.g. *Calb1, Cbln1, Cbln3*), none of which were significantly altered in expression in microglia-deficient cerebellum. **Cluster 4**, enriched in olfactory bulb, included genes for regulators of adult neurogenesis (*Dcx, Dlx1, Dlx2, Dlx5, Tbx21, Sall3, Lgr5, Lgr6, Sp8*) (Tepe et al., 2018) which were expressed equally in WT and *Fireko* mice. Similarly, **Cluster 7** was enriched in striatum and contained multiple genes encoding transcription factors (e.g. *Foxp1, Six3*) required for development of striatal neurons (Yang et al., 2021). There was no significant effect of microglial deficiency on expression of transcripts associated with the vasculature or blood brain barrier (BBB) (e.g. *Ckdn5, Flt1, Vwf, Abcb1, Pecam1*) in any brain region (**Table S1, S2**).

### Microglia deficiency has limited impacts on adult motor function

CRL patients exhibit severely impaired cognitive and motor functions (Konno et al., 2018). Motor function and coordination were assessed in *Fireko*^Edin^ mice using the CatWalk assay (Garrick et al., 2021). Previous analysis of aged *Fireko*^Edin^ mice was based on groups of 4-5 animals (McNamara et al., 2023; Munro et al., 2024). Here we analysed larger groups, and each mouse was tested weekly from 8-40 weeks of age, effectively acting as its own control. On all measured parameters, *Fireko* mice remained indistinguishable from WT and showed no evidence of age-related decline (**Figure S1A**).

### Csf1r expression is restricted to microglia and macrophages

There have been several studies claiming that *Csf1r* is expressed by neurons (Chitu et al., 2022) as well as microglia. We crossed the C57BL/6J *Csf1r*-FusionRed (FRed) knock-in transgenic reporter which reports CSF1R protein synthesis (Grabert et al., 2020) to the widely used *Cx3cr1-*EGFP microglia-specific reporter. In the brains of these mice, there was a complete overlap between *Csf1r-*FRed, *Cx3cr1*-EGFP and IBA1 (**Figure S1B**). Hence, there was no evidence of the reported expression of CSF1R protein in Purkinje cells (Murase and Hayashi, 1998), layer 5 cortical neurons (Clare et al., 2018) or the dentate gyrus in the hippocampus (Luo et al., 2013). The microglia-restricted expression of *Cx3cr1*-EGFP is also incompatible with reported *Cx3cr1* expression in hippocampal neurons (Meucci et al., 2000). Flow cytometry analysis of the two transgenes in isolated microglia and BAM, defined by expression of CD45 and CD11b, is shown in **Figure S1C**. Consistent with their high expression of *Csf1r* mRNA, *Cx3cr1*-EGFP^+^ microglia uniformly express high levels of *Csf1r-*FRed and F4/80. Consistent with previous RNA-seq data (Kiani Shabestari et al., 2022), localization of *Csf1r* mRNA (Kana et al., 2019) and the *Csf1r-*EGFP reporter transgene (Elmore et al., 2014) we conclude that CSF1R is entirely restricted to myeloid cells regardless of mouse genetic background.

### Postnatal survival and hydrocephalus in Fireko mice depends on genetic background

The original *Csf1r*^ΔFIRE^ mutation was introduced in C57BL/6J x CBA/J F1 mice (BCBF1) and crossed to C57BL/6J to generate the 3:1 C57 x CBA background *Fireko*^Edin^ mice (Rojo et al., 2019). The *Csf1r*^ΔFIRE^ allele was crossed 10 times to C57BL/6JCrl in Edinburgh. On this background there was extensive perinatal loss and all *Fireko* mice identified by genotyping at 11 days were euthanized due to hydrocephalus. We imported C57BL/6J *Csf1r*^ΔFIRE/+^ mice to Australia, rederived by crossing to C57Bl/6JArc, then crossed to the *Csf1r*-EGFP reporter line (Sasmono et al., 2003) to generate the congenic *Fireko*^UQ^ line. Surprisingly, although the reporter line was also backcrossed at least 10x to C57BL/6JArc, 15% of pups (33/223) from heterozygous matings were genotyped as *Csf1r*^ΔFIRE/ΔFIRE^ compared to the expected 25% Mendelian frequency. Like *Csf1r^-/-^* mice on the alternative FVB/J background (Elmore et al., 2014) around a third of surviving *Fireko*^UQ^ pups developed severe hydrocephalus (HC) requiring euthanasia by 3-5 weeks. The remainder were long-lived, healthy and fertile.

To understand the nature of HC, Evans Blue dye was injected into the ventricles of affected *Fireko*^UQ^ mice. Severely-affected mice had massively enlarged lateral ventricles (**Figure 2A**) which precluded analysis. In mice with less extreme HC, the lateral ventricles accumulated Evans Blue, whereas the dye was excluded from the third and fourth ventricles and cerebellum (**Figure 2B**) suggesting occlusion of the foramina of Monro, the fine slit-like channel linking the lateral ventricles to the third ventricle. This pattern suggests non-communicating HC. Consistent with this interpretation, *Fireko*^UQ^ mice that did not succumb to early postnatal HC did not show evidence of subsequent ventricular enlargement (**Figure 2C**). To confirm the effect of genetic background on HC in microglia-deficient mice, *Fireko*^UQ^ mice were bred with WT CBA and BALB/c mice, and *Csf1r*^ΔFIRE/+^ F1 progeny were intercrossed to generate F2 mice of all genotypes. On the two mixed backgrounds there was no longer any significant pre-weaning loss. HC was observed in only 3/83 C57BL/6J x CBA and 0/163 C57BL/6J x BALB/c *Fireko* F2 progeny.

**Figure 2.**
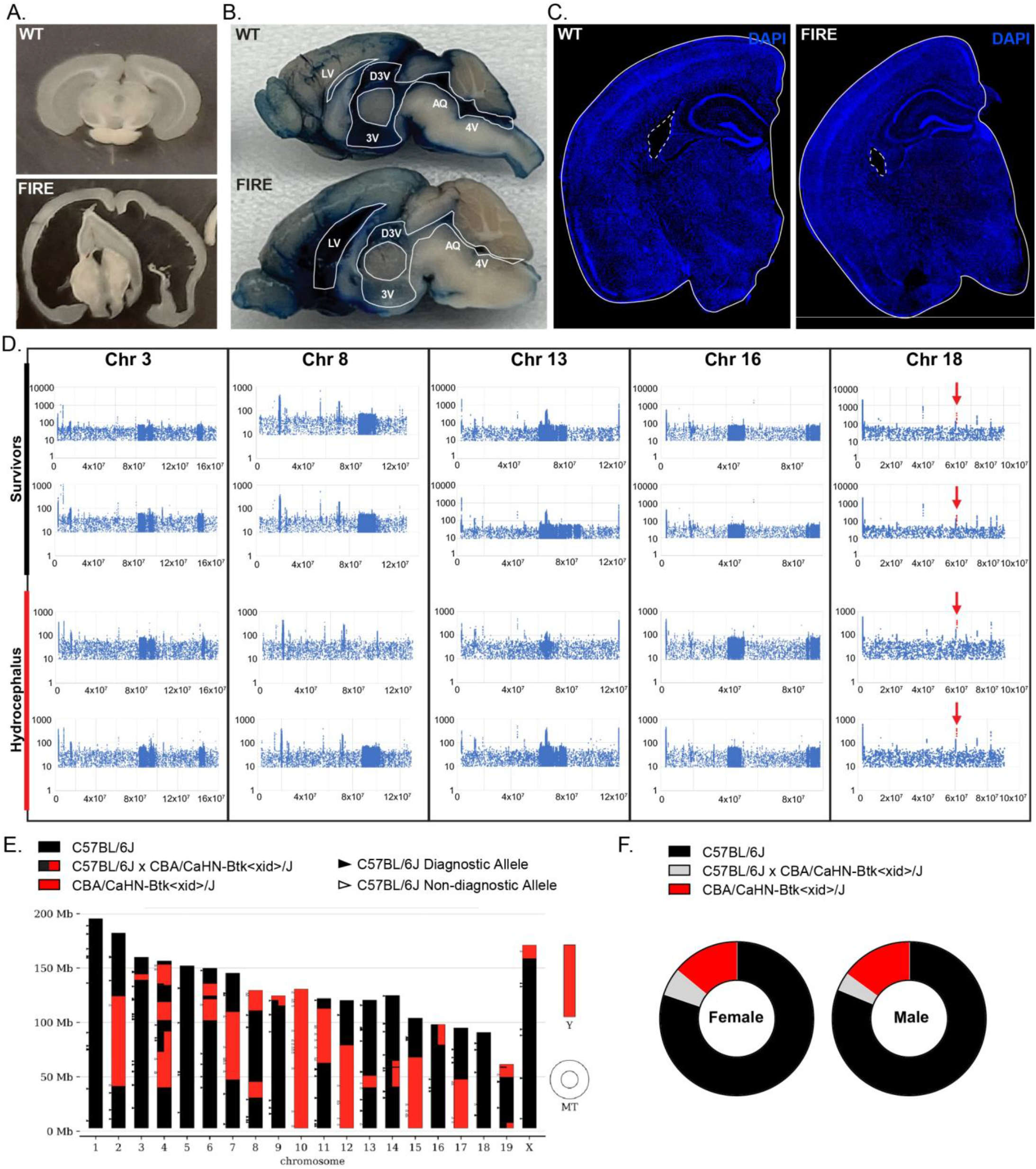
Severe hydrocephalus in *Fireko*^UQ^ mice. (A) Coronal sections of brains from a 5-week-old WT and *Fireko*^UQ^ (FIRE) mouse that developed hydrocephalus requiring euthanasia highlight extreme enlargement of the lateral ventricles. (B) Medial view of the brain of a WT (upper) and a *Fireko*^UQ^ (lower) mouse showing less severe ventricular enlargement. Both mice were injected with Evans Blue dye. Note the dilated lateral ventricle (LV) in the *Fireko* and the exclusion of dye from the third ventricle and dorsal third ventricle (3V, D3V), aqueduct (AQ), fourth ventricle (4V) and hindbrain compared to the WT. (C) Coronal sections of typical 30-week-old WT and *Fireko*^UQ^ (FIRE) brains stained with DAPI. The lateral ventricles are highlighted. (D) DNA was isolated from mutant mice that developed hydrocephalus and long-term survivors on the congenic C57BL/6J *Fireko*^UQ^ background. Whole genome sequences were generated from two pools from each group, each containing DNA from three or four mice. Sequences were mapped to the C57BL/6J reference genome. Images show the read count for non-reference sequences for each nucleotide. Regions derived from non-C57BL/6J origin are shown by dense incidence of non-reference variants, highlighting blocks detected specifically on Chr 3, 8, 13 and 16. Datapoints in red (arrow) on Chr 18 highlight an excess of non-reference reads mapped to the location of the *Csf1r* gene. The promoter used in the *Csf1r*-EGFP reporter is from a non-C57BL/6J source and contains a mutated start codon, accounting for these reads. X-axis – genomic position on chromosome; Y-axis – Log_10_ number of reads over position. (E-F) DNA was isolated from 2 male and 2 female *Fireko*^Edin^ mice and analysed using the MiniMUGA mouse genotyping array. The typical chromosome map (E) shared by the 4 mice and averaged data (F) are consistent with the predicted 25% contribution of CBA/J to the genetic background, occurring primarily as blocks of homozygosity.

We speculated that *Fireko*^UQ^ mice that did not develop HC might be enriched for residual non-C57BL/6J alleles that mitigate perinatal loss or the development of HC. To test this possibility, we performed whole genome sequencing on two separate pools of *Fireko*^UQ^ mice with HC and long-term survivors that did not develop HC (**Figure 2D**). The four pools were not distinguished from each other based on genomic sequence. Pools shared non-reference single nucleotide variants throughout the genome as well as small blocks of linkage disequilibrium; defined regions with a high frequency of alternate alleles (compared to the C57BL/6J reference) on chromosomes 3, 8, 13 and 16. One of two separate blocks on Chr 16 was identified as the likely location of the *Csf1r*-EGFP transgene. An apparent breakpoint at Chr16:91,207,427 joins to the *Csf1r* promoter sequence included in the transgene and a second at Chr16:91,183,085 (GRCm38) joins to the vector sequence. *Inter alia,* the genomic DNA sequence revealed that the *Csf1r*^ΔFIRE^ allele was introduced on the C57BL/6J Chr 18 in the F1 founder since there was no enrichment for non-reference sequences in the vicinity of the *Csf1r* locus (red arrow). A single peak of non-reference sequence on Chr 18 is a consequence of mapping 10-20 copies of the 7.2kb *Csf1r* promoter sequence contained within the multi-copy *Csf1r-*EGFP transgene to the reference sequence that lacks the transgene. These data confirm that despite extensive backcross, the *Csf1r*-EGFP mice have contributed segments of non-reference C57BL/6J genomic DNA to the *Fireko*^UQ^ line.

To explore the genetic basis for the difference in HC incidence between *Fireko*^UQ^ and *Fireko*^Edin^ we performed SNP genotyping. As shown in **Figure 2E-F**, reflecting brother-sister mating of the founders and subsequent inbreeding, the *Fireko*^Edin^ line is effectively a recombinant inbred line with most of the genome in all animals genotyped homozygous for either C57BL/6 or CBA SNPs. The regions of homozygous C57BL/6J background likely contribute to perinatal lethality whilst those of CBA origin, which include the MHC locus on Chr 17, mitigate against HC. Taken together, the genetic analysis indicates that postnatal lethality and HC in *Fireko* mice are independent probabilistic complex traits with increased likelihood on the C57BL/6J background.

### Development of brain-associated macrophages in Fireko mice is not affected by genetic background

Adult *Fireko*^Edin^ mice lack microglia but retain BAM (Rojo et al., 2019) as confirmed in the RNA-seq analysis above. **Figure 3A** shows immunolocalisation of IBA1 and the microglia-restricted TMEM119 marker in the hippocampus of adult (15 week) *Fireko*^UQ^ mice. IBA1^+^/TMEM119^-^ BAM were detected in association with blood vessels identified by their associated GFAP-expressing astrocytes. IBA1 staining of coronal sections through the brain of *Fireko*^UQ^ neonates confirmed the complete absence of parenchymal microglia at birth (**Figure 3B**). IBA1^+^ cells on meningeal surfaces and lining blood vessels were also less abundant at this age compared to WT controls (**Figure 3B**). Epiplexus macrophages, the supraependymal population within the choroid plexus, detected in WT using IBA1 staining (Yellow arrowheads; **Figure 3C**), were absent in the *Fireko*^UQ^ neonates. By contrast, stromal choroid plexus macrophages were present but reduced compared to WT mice (White arrowheads, **Figure 3C**). **Figure 3C** also shows localisation of GFAP, which is expressed by ependymal cells in neonatal mice (Wu et al., 2020). In both WT and *Fireko*^UQ^ neonates, GFAP^+^ cells formed a continuous lining of the ventricles. In terms of brain myeloid populations in neonates, the *Fireko*^UQ^ mirror the pattern reported previously (Munro et al., 2020). We conclude that the development of HC in congenic *Fireko*^UQ^ compared to *Fireko*^Edin^ mice is not associated with a more penetrant loss of myeloid cells.

**Figure 3.**
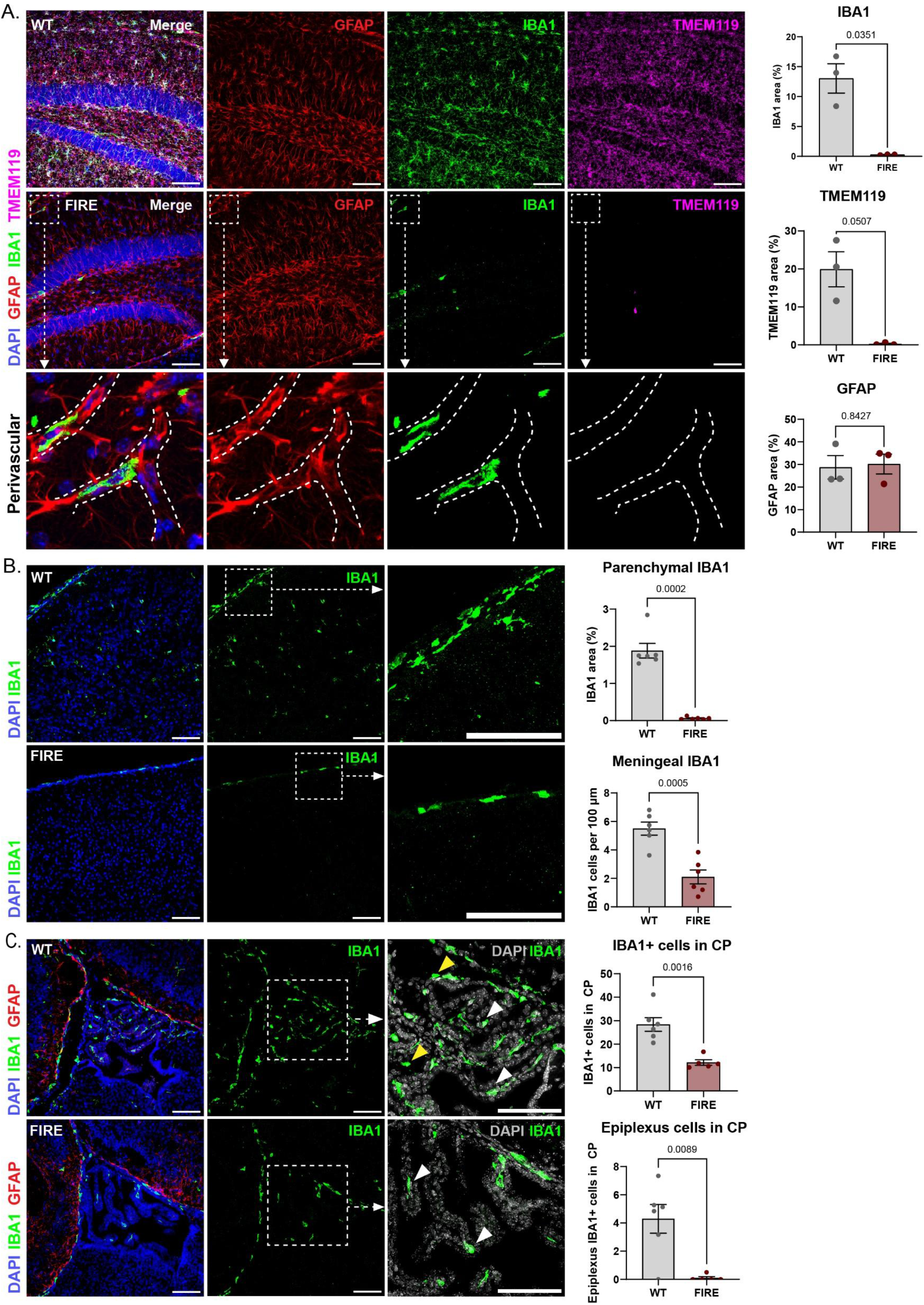
Selective loss of microglia in adult and neonatal *Fireko*^UQ^ mice. (A) Free-floating sections from adult WT or *Fireko*^UQ^ (FIRE) hippocampus were stained to detect GFAP, IBA1, and TMEM119 and counter-stained with DAPI as indicated. Representative images are shown. Percentage of positive staining is shown quantified to the right, from three individual animals per genotype. Differences between groups were analysed by unpaired t test with Welch’s correction. P values are displayed on each graph. Lower panels show boxed region at higher resolution to highlight retention of IBA1^+^ perivascular macrophages in *Fireko* mice. Scale bars = 100 µm. (B-C) Cryosections of cortex (B) and choroid plexus (C) from neonatal WT or *Fireko*^UQ^ (FIRE) mice were stained to detect IBA1 and GFAP as indicated. Percentage of positive IBA1 staining is shown quantified to the right, from at least 6 different regions across brain parenchyma. The number of IBA1^+^ meningeal macrophages per 100 µm is also shown to the right, quantified from at least 5 different regions of cortex across both genotypes. In representative images of the choroid plexus (C), white arrows indicate stromal macrophages. Epiplexus macrophages indicated by yellow arrows in WT were not detected in *Fireko* mice. The number of IBA1^+^ cells and epiplexus macrophages was quantified from at least 5 different regions from 2-3 animals per genotype. Scale bars = 100 µm. Graphs show mean ± SEM, analysed by unpaired t test with Welch’s correction. P values are reported on each graph.

### Analysis of Fireko mice on a C57BL/6J x CBA/J F2 background

In view of the 100% survival and the absence of HC in *Fireko*^UQ^ mice on the C57BL/6J x CBA/J F2 background, we considered whether neuropathology reported in *Fireko*^Edin^ mice (Munro et al., 2020) might be associated with the regions of C57BL/6 or CBA homozygosity. We re-analysed the phenotype on the F2 background which we refer to as *Fireko*^BCBF2^. We first conducted standard behavioral assays at 30 weeks of age. We found no significant differences between WT and *Fireko* mice in novel object recognition, the marble burying test, or elevated plus maze, commonly applied tests of memory and anxiety-like behavior (Chen et al., 2024) (**Figure 4A-C**). We also performed Magnetic Resonance Imaging (MRI) experiments including susceptibility weighted imaging (SWI) and diffusion tensor imaging (DTI) on these mice. SWI is commonly used to detect cerebral microbleeds, whereas DTI detects white matter pathology (Althobity et al., 2023) and is commonly applied in diagnosis of CRL (Konno et al., 2018). We found no evidence of hyperintense deep white matter diffusion dots, cortical atrophy or ventricular enlargement typical of human pathology in any genotype (**Figure 4D**). Thinning of the corpus callosum is commonly detected in CRL patients (Konno et al., 2018) but we saw no difference in the DTI imaging of the corpus callosum, anterior commissure, and hippocampal commissure (**Figure 4E**). Quantitative analysis showed a slight decrease in the fractional anisotropy (FA) intensity in the hippocampal commissure of both *Csf1r*^ΔFIRE/+^ and *Fireko* mice compared to WT, but no significant changes in FA volume were detected in any region **(Figure S2A-B**). Consistent with this view, quantification of myelin basic protein (MBP) staining did not indicate significant demyelination or hypermyelination in the corpus callosum, cortex, or striatum in the *Fireko*^BCBF2^ mice at this age (**Figure 4F, Figure S2C**).

**Figure 4.**
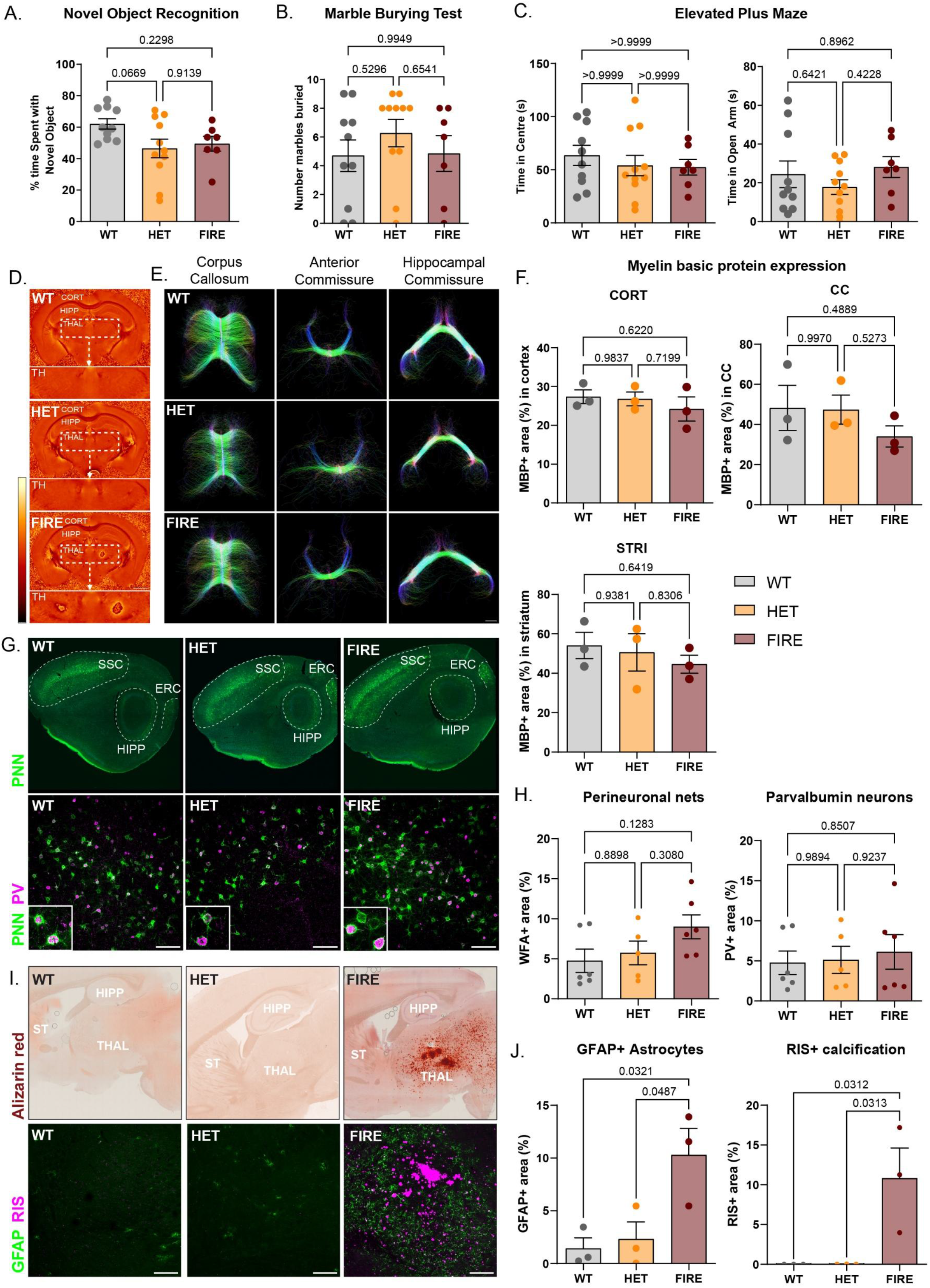
Functional and histological analysis of *Fireko*^BCBF2^ mice. Standard behavioural tests; (A) Novel object recognition. (B) Marble burying test. (C) Elevated plus maze were performed on cohorts of 30-week-old WT and *Fireko*^BCBF2^ mice on the C57BL/6J x CBA F2 mice as described in Methods. Graphs show mean ± SEM of at least seven mice per genotype. Data were analysed using an ordinary one-way ANOVA with Tukey’s multiple comparisons test. (D) Representative images of susceptibility weighted imaging (SWI) showing cortex (CORT), hippocampus (HIPP), and thalamus (THAL) in 30-week-old WT, *Csf1r*^ΔFIRE/+^ (HET) and *Csf1r*^ΔFIRE/ΔFIRE^ (FIRE) C57BL/6J x CBA F2 mice. Boxed region highlights bilateral hypointensity in the thalamus of *Fireko* mice. (E) Representative images of diffusion tensor imaging (DTI) of corpus callosum, anterior commissure and hippocampal commissure of the same 30-week-old mice. Scale bar = 50 mm. (F) Brains of 30-week-old mice were stained with myelin basic protein (MBP; **Figure S2C**) and percentage area of MBP^+^ staining in cortex (CORT), corpus callosum (CC), and striatum (STRI) was quantified. Graphs show mean ± SEM of three mice per genotype. Data were analysed using an ordinary one-way ANOVA with Tukey’s multiple comparisons test. (G) Brains of 30-week-old mice were stained with *Wisteria floribunda* agglutinin (WFA) to detect perineuronal nets (PNN), and antibody against parvalbumin neurons (PV). PNNs were detected in the somatosensory cortex (SSC), the entorhinal cortex (ERC) and the dentate gyrus of the hippocampus (HIPP). Lower panels show detection of both markers in the SSC at higher resolution. Scale bar = 100 μm. (H) Graphs show mean ± SEM area of WFA+ and PV+ staining of two quantified regions per mouse, from three mice per genotype. (I) Upper panel shows representative images of brains from same 30-week-old mice stained with Alizarin Red. Lower panels show representative images of the thalamus stained with GFAP and AF647-RIS. Percentage area quantification is shown on the right (J); graphs show mean ± SEM from three animals per genotype, analysed using ordinary one-way ANOVA followed by Tukey’s multiple comparison test. All P values are reported on each graph.

Perineuronal nets (PNN) are brain extracellular matrix structures that regulate synaptic functions and protect neurons from stress (Testa et al., 2019). Microglial depletion was previously found to increase PNN density whereas microgliosis associated with heterozygous *Csf1r* knockout mutation on the C57BL/6J background led to PNN depletion (Arreola et al., 2021; Barahona et al., 2025; Liu et al., 2021; Valdearcos et al., 2025). In the cortex, PNN are commonly associated with inhibitory parvalbumin (PV) interneurons (Reichelt et al., 2019). They can be detected by staining with the lectin, *Wisteria floribunda* agglutinin (WFA) (Arreola et al., 2021). Consistent with published reports, we detected WFA staining in the somatosensory cortex (SSC), entorhinal cortex (ERC), and hippocampus (HIPP) often surrounding PV^+^ neurons (**Figure 4G**). In the *Fireko*^BCBF2^ line, there was no significant difference in the localization of PNN or PV interneurons, or staining intensity between WT, *Csf1r*^ΔFIRE*/+*^ and *Fireko* mice at 30 weeks of age (**Figure 4H**). Brain calcification is a cardinal feature of both dominant and recessive CRL and precedes detectable white matter degeneration (Konno et al., 2017). Extensive age-dependent astrocytosis and calcification of the thalamus is the most notable reported brain pathology in *Fireko* mice (Bradford et al., 2022; Chadarevian et al., 2024; Kiani Shabestari et al., 2022; Munro et al., 2024). Re-analysis of the uninfected control cohort from our prion disease study using *Fireko*^Edin^ mice (Bradford et al., 2022) revealed the accumulation of GFAP-positive astrocytes in the thalamus in a subset of *Fireko* mice as early as three months of age, before any calcification was detectable by Alizarin Red staining (**Figure S3A**).

A detailed analysis of spontaneous brain lesions in C57BL/6J mice revealed prominent thalamic neuronal inclusions at a relatively young age (Tarrant et al., 2020). We therefore considered the possibility that thalamic pathology depends upon the predominant C57BL/6J background in *Fireko*^Edin^ mice. However, a notable feature of the SWI in **Figure 4D** is the detection of hypointensities in the thalamus of *Fireko*^BCBF2^ mice. Extensive calcification was confirmed by Alizarin Red and AF647-risedronate (RIS) staining in the thalamus of these 30-week-old *Fireko* mice, closely associated with accumulation of GFAP^+^ astrocytes (**Figure 4I, J**).

### Sustained repopulation of vacant tissue macrophage territories in Fireko mice without donor contribution to blood monocytes

We reported elsewhere that IP BMT of WT cells from a *Csf1r*-FusionRed (FRed) donor repopulated all the vacant macrophage territories in *Fireko*^UQ^ mice by 12 weeks post transfer without giving rise to circulating monocytes (Liu et al., 2026). Here, we asked whether repopulation was sustained with 27 weeks post transfer. The F4/80^high^ large peritoneal macrophage (LPM) population is progressively replaced by recruited monocytes which give rise to F4/80^low^ small peritoneal macrophage (SPM) precursors (Bain et al., 2016). In the cavity of *Fireko*^UQ^ BMT recipients F4/80^high^ LPM were fully restored and uniformly *Csf1r-* FRed^+^ (**Figure S4A**). Consistent with analysis 12 weeks post BMT, where FRed was not detected in CD11b^+^ cells in bone marrow, blood or spleen (Liu et al., 2026), in the blood of BMT recipients 27 weeks post-transplant, CSF1R remained selectively reduced in Ly6C^high^ monocytes and *Csf1r-*FRed^+^ was not detectable (**Figure S4B**). Whole mount images in **Figure S4C** confirm that repopulation of various vacant territories in the *Fireko* BMT recipients was sustained 27 weeks post BMT. Immunostaining of brains stained with IBA1 also confirm the repopulation of brain parenchyma with IBA1^+^ FRed^+^ cells in *Fireko* BMT recipients, and that FRed^+^ cells were not present in the parenchyma of WT BMT recipients, highlighting the requirement of a vacant niche for repopulation to occur (**Figure S4D**).

### Repopulation of the brain with bone marrow-derived microglia-like cells mitigates the development of neuropathology

We next asked whether age-dependent neuropathology was evident in *Fireko*^UQ^ mice and whether it was prevented or potentially exacerbated by bone marrow-derived cells. Consistent with evidence from the *Fireko*^BCBF2^ line, we detected increased GFAP^+^ astrocytes in the thalamus of *Fireko*^UQ^ mice at 15 weeks, prior to the development of extensive calcium deposits at 30 weeks (**Figure S3B, C; Figure 5D**). The brain (**Figure 5A-C, Figure S5A)** and spinal cord (**Figure S5B, C**) of *Fireko*^UQ^ BMT recipients 27 weeks post transfer were completely repopulated with IBA1^+^ microglia-like cells of donor origin, displaying the regular tiling that is characteristic of microglia but with a somewhat less ramified morphology. In the BMT recipients, astrocytes detected by GFAP staining were not significantly elevated compared to WT and calcification observed in age-matched *Fireko* controls was absent (**Figure 5D-F**). As reported previously in *Fireko*^Edin^ mice (Kiani Shabestari et al., 2022; Munro et al., 2024), we also detected SMI315^+^ axonal spheroids that were prevented by BMT (**Figure S3D**).

**Figure 5.**
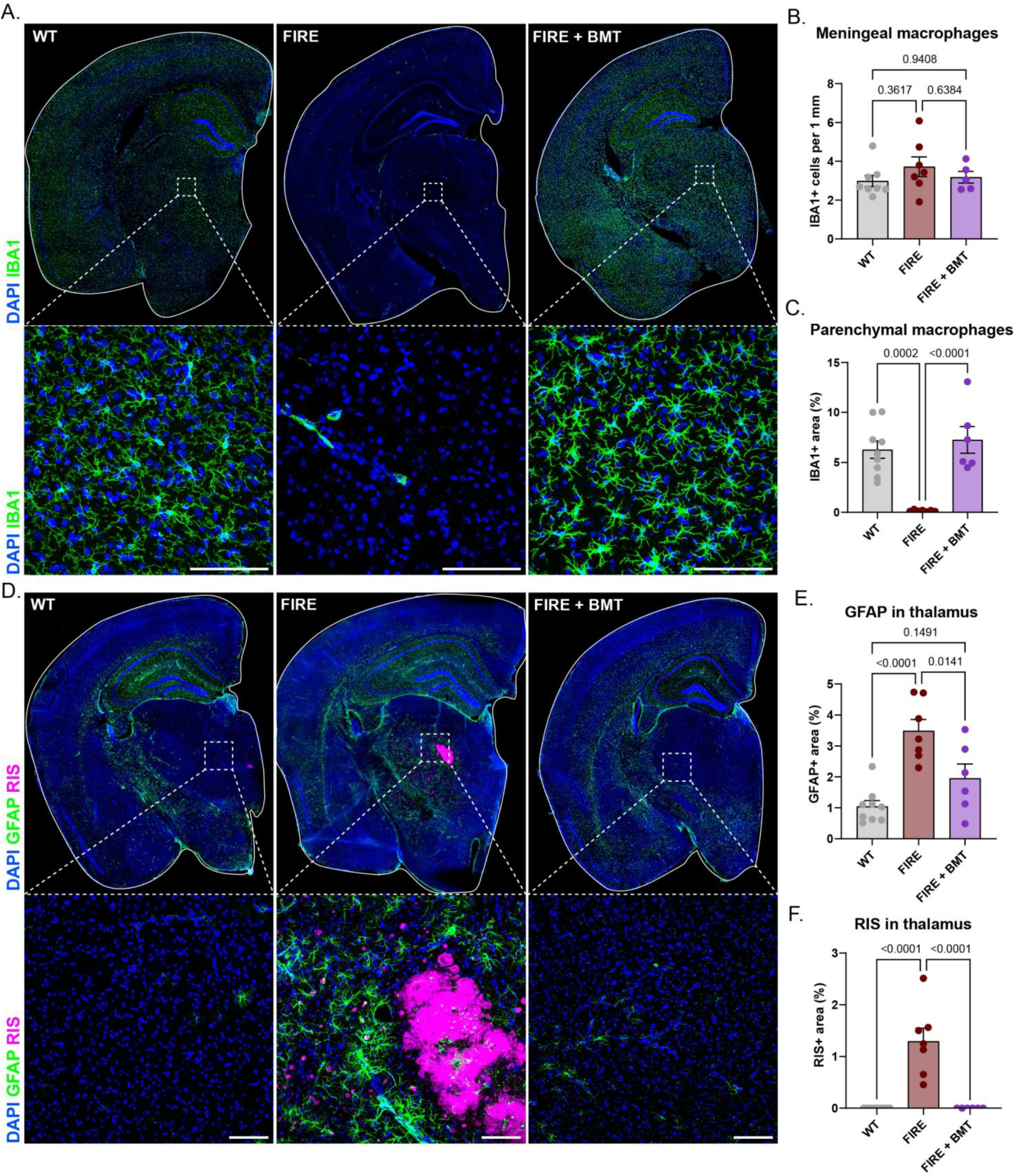
Repopulation of the brain following WT bone marrow cell transfer prevents thalamic calcification in *Fireko*^UQ^ mice. Three-week-old *Fireko*^UQ^ mice received WT bone marrow cells by i.p. injection (BMT). Controls were WT mice or untreated *Fireko* mice of the same age. (A) Free-floating coronal sections from 30-week-old WT, *Fireko* (FIRE), or *Fireko* + BMT (FIRE + BMT) mice were stained to detect IBA1 (A-C), or GFAP and AF647-RIS^+^ calcification (D-F) as indicated. Representative images are shown; lower panels show highlighted region at higher resolution. Scale bars = 100 µm. (B) The number of IBA1^+^ meningeal macrophages per 1 mm, quantified from at least five mice per genotype. (C) Percentage of IBA1^+^ staining through entire brain hemisphere was quantified from at least six mice per genotype. (E-F) Percentage of GFAP^+^ (E) or RIS^+^ (F) staining in thalamus was quantified from at least six mice per genotype. Graphs show mean ± SEM, analysed by ordinary one-way ANOVA followed by Tukey’s multiple comparisons test. P values are shown on each graph.

To further analyse the impact of BMT in *Fireko*^UQ^ mice, we performed bulk RNA-seq profiling of thalamus and cortex of WT, *Fireko* and *Fireko*+BMT mice. The complete datasets are provided in **Table S4/S5**. **Figure 6** shows differential gene expression analysis between the three states for thalamus (**Figure 6A-C**) and cortex (**Figure 6D-F**). To deconvolute the data, we again performed co-expression analysis on the combined dataset. The outcomes are provided in **Table S7.** Several co-expression clusters were associated with genotype or repopulation following BMT. **Cluster 8** contains transcripts that were detected at higher levels in BMT recipients compared to age-matched WT controls and includes markers associated with microglia-like BM-derived cells in previous studies (e.g. *Clec7a, Clec12a*, *Cst7, Lgals3, Itgax, Mrc1, Ms4a7, Msr1*) (Bastos et al., 2025; Bennett et al., 2018; Du et al., 2026; Shemer et al., 2018). Transcripts in **Cluster 9** and **45,** including microglial markers *Aif1* (IBA1) and *Cx3cr1,* shared a pattern of reduced detection in the *Fireko* and restoration in BMT recipients. Transcripts most strongly correlated with *Csf1r,* including *P2ry12,* form the smaller **Cluster 80,** almost undetectable in *Fireko* and incompletely restored following BMT. Genes of interest from these four clusters are listed below each graph in **Figure 6G**. *Gfap, C4b* and *Serpina3n* were each increased in both brain regions in *Fireko* mice (**Figure 6H**). This response was neither prevented nor exacerbated by BMT. Transcripts associated with oligodendrocytes and myelination did not form a separate cluster. Considering these genes separately, the expression of the transcriptional regulator, *Myrf*, and target genes (*Mbp, Mag, Plp1*) in the thalamus of BMT recipients was not significantly different between WT and *Fireko* and was not altered by BMT (**Figure 6I**). In scRNA-seq data, Munro *et al*. (Munro et al., 2024) identified a set of transcripts in addition to *C4b* and *Serpina3n* that were increased in oligodendrocytes from *Fireko* brains. From this set, we detected increases in *Flnc, Emp3, Bcl3, Trf* in our 30-week-old *Fireko* mice (**Figure 6J**). There was no consistent effect of BMT on their expression. These authors also reported an apparent reduction in NeuN^+^ neurons in the thalamus of *Fireko* mice. However, we found no significant reduction in *Foxred1* (encoding NeuN), nor in any of the markers that define individual thalamic nuclei in the mouse (*Tcf7l2, Gbx2, Lef1, Prox1, Six3, Esrrg, Rora, Gad2,* (Roy et al., 2022)) (**Table S4**).

**Figure 6.**
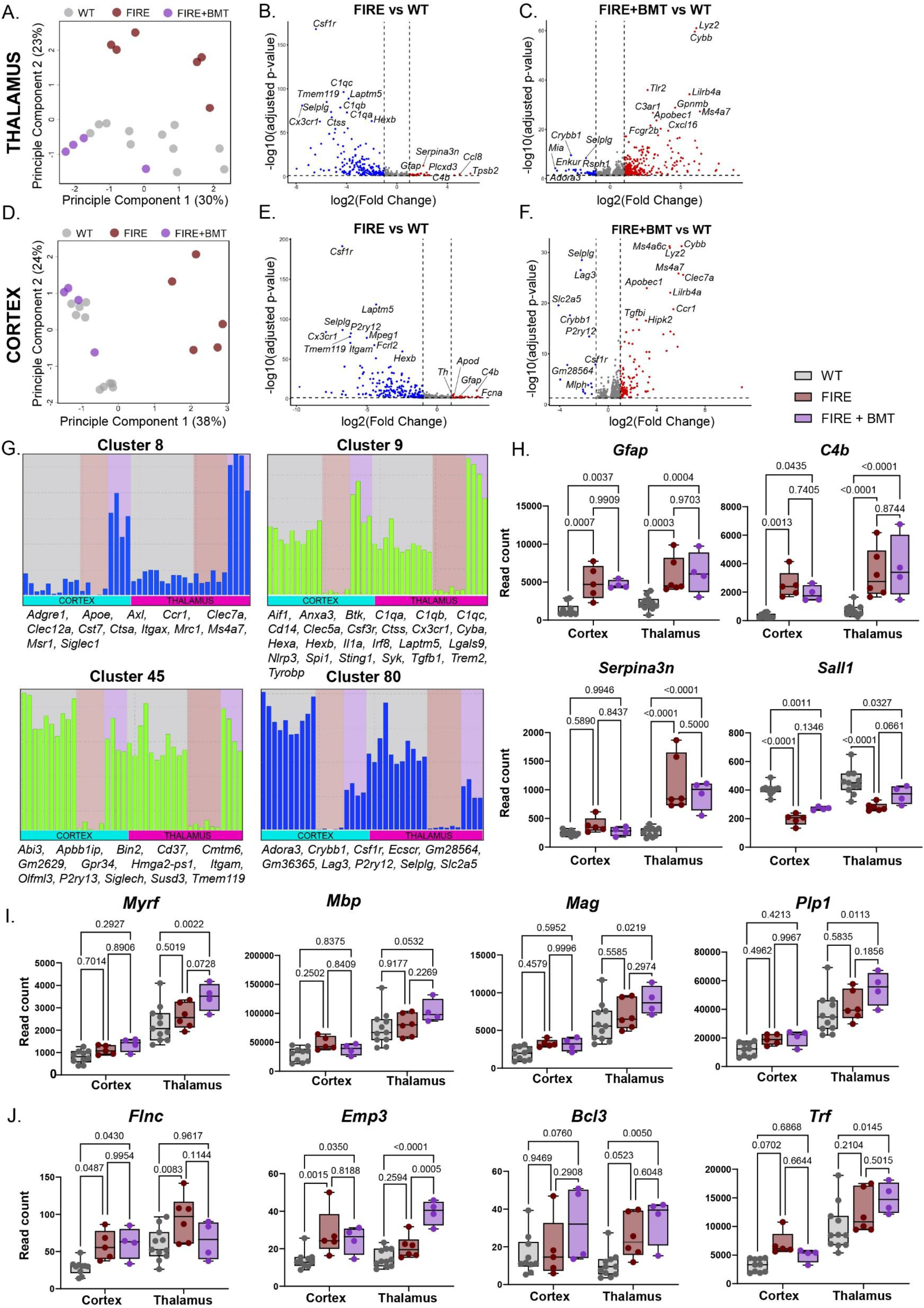
Transcriptomic analysis of the impact of wild-type bone marrow cell transfer on thalamus and cortex of *Fireko*^UQ^ mice. Thalamus and cortex from 30-week-old *Fireko*^UQ^ mice that received BMT, with WT and untreated *Fireko* controls, were dissected and analysed by total RNA-seq. (A-F) Principal component analysis and volcano plots of differentially expressed genes in thalamus (A-C) and cortex (D-F) are shown. Genes significantly upregulated in *Fireko* (B, E) or *Fireko* + BMT (C, F) compared to WT brains are shown in red, while downregulated genes are blue. (G) The complete RNA-seq dataset was clustered using *Graphia,* and the shared profile of four clusters (8, 9, 45, and 80) associated with *Fireko* genotype and the effect of BMT are shown with representative genes of interest listed below. (H-J) RNA expression of markers of neuropathology (H), microglia (*Sall1*) and myelination (I), and oligodendrocytes (J) are graphed. Each point is an individual animal. Graphs show mean ± SEM, analysed by ordinary two-way ANOVA. Tukey’s multiple comparisons test was used to compare genotypes within each brain region (simple effects within columns). P values are shown on each graph.

We performed quantitative analysis of IBA1 staining on multiple brain regions comparing BMT recipients and their age matched WT controls. The percentage area positive for IBA1 was slightly lower in the cortex of BMT recipients compared to WT but the number of IBA1^+^ cell bodies was elevated in thalamus and corpus callosum (**Figure 7A-B**). The area of each IBA1^+^ cell was also increased in BMT recipients compared to WT (**Figure 7C**). Immunolocalisation of F4/80 provided the most striking marker of the distinct phenotype of bone marrow-derived cells, highlighting both the large increase in fluorescence intensity and altered morphology compared to WT microglia throughout the brain (**Figure 7D, E**).

**Figure 7.**
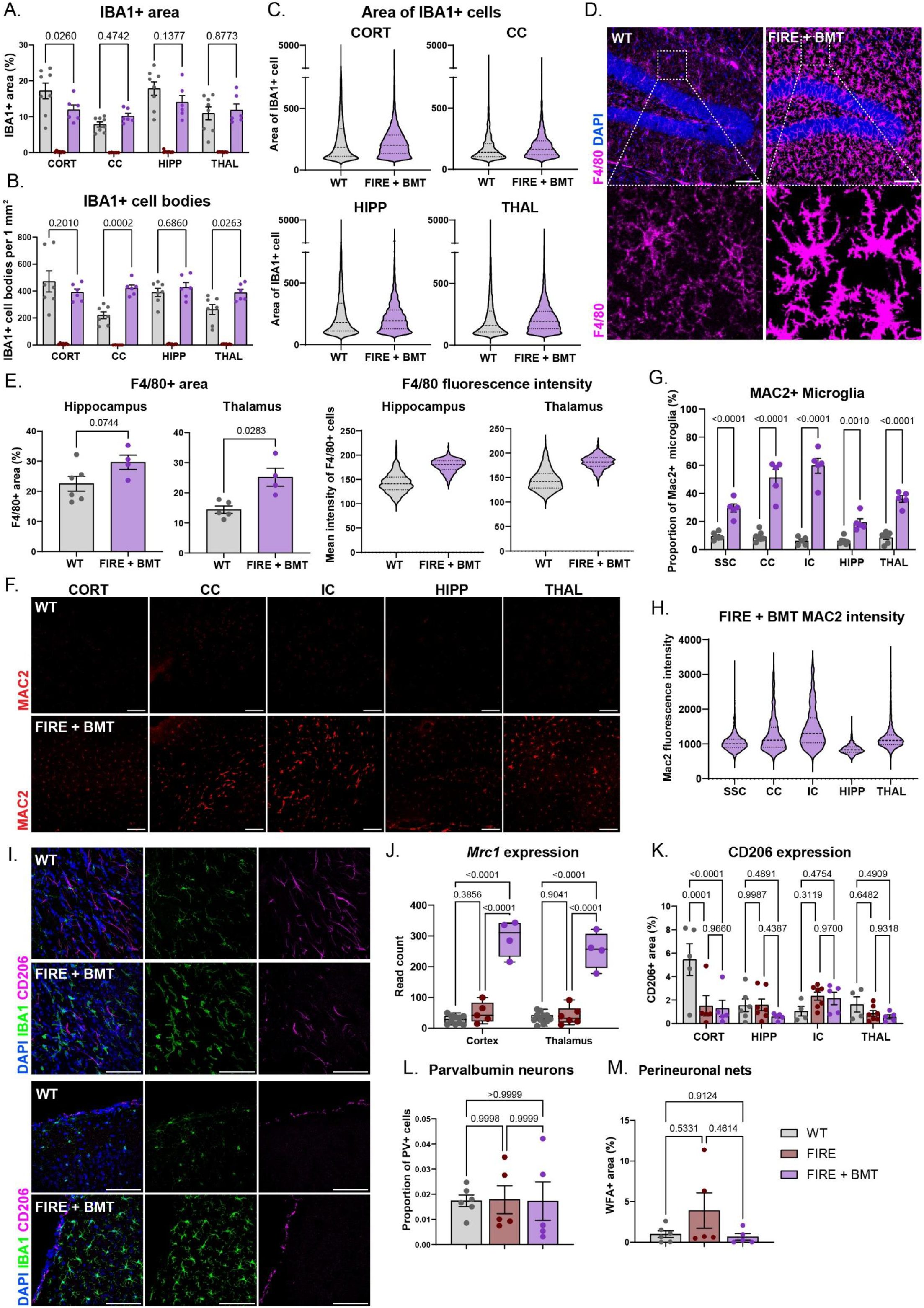
Phenotype analysis of microglia-like cells derived from wild-type bone marrow in *Fireko* mice. (A) Percentage area of total IBA1^+^ staining was quantified from cortex (CORT), corpus callosum (CC), hippocampus (HIPP), and thalamus (THAL) from 30-week-old WT, *Fireko*^UQ^ (FIRE), and *Fireko*^UQ^ + BMT (FIRE + BMT) mice. (B) The number of IBA1^+^ cells was quantified for each region, from at least six mice per genotype. Bar graphs show mean ± SEM, analysed by ordinary two-way ANOVA followed by Tukey’s multiple comparisons test. (C) Violin plots show the size of each IBA1^+^ cells in each region, calculated using QuPath as described in methods. Data from at least five mice per genotype were pooled. (D) Free-floating coronal sections from the same mice were stained for F4/80. (E) Percentage area and fluorescence intensity of F4/80^+^ stain in hippocampus and thalamus were quantified. P values are shown on each graph. (F-H) Free-floating sections of the same brains were stained to detect MAC2 (LGALS3). Representative images of regions including internal capsule (IC) are shown. Proportion of MAC2^+^ microglia (G) and mean fluorescence intensity (H) are shown quantified to the right. (I-K) Free-floating sections were stained to co-localise IBA1 and CD206 in thalamus (top) and cortex (bottom). RNA expression of *Mrc1* is shown to the right (J), compared to CD206 expression quantified from images (K). Sections were also stained to detect parvalbumin neurons and perineuronal nets (images not shown). Proportion of PV^+^ cells (L) and percentage area of WFA^+^ stain (M) are shown. Data in (E, G) are mean ± SEM of at least four mice per genotype, analysed by unpaired t test with Welch’s correction. All other bar graphs show mean ± SEM of at least four mice per genotype, analysed by ordinary two-way ANOVA followed by Tukey’s multiple comparisons test. P values are shown on each graph. All scale bars = 100 μm.

Whereas F4/80 appeared relatively uniform and coincident with IBA1, MAC2 (*Lgals3*) was more variable; the strongest expression was on microglia located within white matter (corpus callosum, internal capsule), and expression was lower in hippocampus, cortex, and thalamus suggesting a path of repopulation migrating along white matter tracts (**Figure 7F-H).** Finally, although *Mrc1* mRNA was also increased in *Fireko*+BMT compared to WT, CD206 immunoreactivity was restricted to border-associated macrophage populations and IBA1-negative cells (possibly astrocytes (Burudi et al., 1999)), and protein expression was not significantly increased in BMT recipients (**Figure 7I-K**). As discussed above, PNN associated with PV neurons were reported to be sensitive markers of microglial dysregulation. However, these markers did not distinguish WT, *Fireko* and *Fireko+*BMT mice (**Figure 7L, M**).

## DISCUSSION

### Compensatory changes in gene expression in microglia-deficient mice

Compared to related cells of the mononuclear phagocyte system, microglia have a distinct gene expression profile with selected genes highly over-expressed relative to other tissue resident macrophages (Summers et al., 2020). The CSF1R-dependent set of 250 transcripts identified herein (**Cluster 11** in juveniles; **Clusters 8/9/45/80** at 30 weeks) includes many known microglia-specific genes (*Crybb1, Cx3cr1, Hexb, P2ry12, Tmem119, Trem2*) and transcriptional regulators (*Spi1* (PU.1), *Irf8, Irf5, Ikzf1* and *Runx1*). One notable feature is the CSF1R-dependent expression of a set of transcripts associated with the generation of vasoactive arachidonic acid metabolites, including *Lpcat2, Plcg2, Alox5, Alox5ap, Ptgs1, Ltc4s*, *Hpgd* and *Hpgds*, that may contribute to the reported regulation of cerebral vascular flow by microglia (Csaszar et al., 2022).

Network analysis of bulk RNA-seq data complements previous single cell RNA-seq data where the expression profile of individual defined neuronal cell types was not dependent upon *Csf1r* genotype (O’Keeffe et al., 2024). The lack of genotype effect on cerebellum-specific genes (**Cluster 1)** contradicts the view that microglia are essential for development of the cerebellum (Kana et al., 2019) and the longitudinal Catwalk analysis in **Figure S1A** did not replicate the motor deficiencies reported in that study attributed to cerebellar dysfunction in the absence of microglia. Similarly, **Cluster 4** and **Cluster 7** argue against an essential regulatory function of microglia in neurogenesis in the subventricular zone (Morton et al., 2018; Ribeiro Xavier et al., 2015) or cell-type specification in the striatum (Knowles et al., 2021), respectively.

In both juveniles and adults, many transcripts considered microglial markers were surprisingly unaffected by microglial deficiency. *Tspo,* encoding the mitochondrial target used to detect microgliosis associated with human neuropathology by PET imaging (Fairley et al., 2023), was unchanged in the microglia-deficient mice. *Nrros,* a leukocyte-specific transcript highly expressed in isolated microglia (Summers et al., 2020; Van Hove et al., 2019) was also unchanged in *Fireko* mice and similarly unaffected in brains of *Csf1r*^-/-^ rats (Patkar et al., 2021). *Nrros*^-/-^ mice develop abnormal microglia, astrocytosis and age-dependent motor deficiencies (Wong et al., 2017). Presumably in the absence of microglia, the *Nrros* gene is upregulated in other cell types, possibly astrocytes (Bedolla et al., 2024; Hasel et al., 2021; Munro et al., 2024). Similarly, microglia in the brain are the main source of insulin-like growth factor 1 (IGF1) which is essential for postnatal neurodevelopment (Hammond et al., 2019; Rusin et al., 2024) but *Igf1* mRNA was unchanged in any brain region in *Fireko* mice (or *Csf1r^-/-^* rats (Patkar et al., 2021)) compared to WT at either time point. Development and survival of layer 5 cortical neurons was reported to be dependent on microglial IGF1 (Ueno et al., 2013) but markers of this population (*Bcl11b, Satb2*) were unaffected in either juveniles or in 30 week cortex (**Table S5**).

*Hexb* is another microglia-enriched gene that distinguishes microglia from BAM in mice (Masuda et al., 2020). *Hexb* knockout mice exhibit many of the neuropathologies of the human HEXB deficiency (Sandhoff disease) (Sango et al., 1996; Sango et al., 1995). Two recent studies showed that microglial replacement by bone marrow transplantation was sufficient to mitigate the pathology in *Hexb*^-/-^ mice (Frosch et al., 2025; Tsourmas et al., 2025). In juvenile and adult *Fireko* mice, *Hexb* mRNA was still detected, albeit reduced to around 10-20% of WT in all brain regions (**Table S1/S2; Table S4/S5**), whereas *Hexa* encoding the other subunit of hexosaminidase was unaffected. Other lysosomal enzymes (e.g. *Dnase2a, Gla, Gba, Galc, Lipa, Gusb, Naga, Naglu, Sumf1*), lysosomal structural genes (Schroder et al., 2010), each associated with a rare lysosomal storage disease (Akhter et al., 2019) and progranulin (*Grn*) which regulates lysosomal function (Tejwani et al., 2026) are selectively expressed in microglia at levels at least 10-fold higher than total brain or other brain cell types (Munro et al., 2024; Zhang et al., 2014). Yet, of all lysosome-related transcripts aside from *Hexb,* only *Gusb, Dnase2a, Manb1* and *Arsb* were significantly reduced in microglia-deficient mice (**Tables S1/S2**) suggesting they are also up-regulated in other cells to compensate. We re-examined scRNA-seq data generated from hippocampus of juvenile WT and *Fireko* mice (Munro et al., 2024). Microglia-enriched transcripts including *Arsb, Man2b1, Hexb, Nrros, Gusb, Ctsd, Grn, Dnase2a*) in this dataset are detectable in both astrocytes and oligodendrocytes. *Dnase2a* is of particular interest as it is required for degradation of nuclear DNA of ingested apoptotic cells. In addition to *Dnase2a,* the phagocytic receptors *Mertk* and *Axl* and ligands *Gas6* and *Pros1* are expressed in both microglia and astrocytes (Damisah et al., 2020; Munro et al., 2024). In bulk RNA-seq, their expression was unchanged in any brain region at any time in the *Fireko*. These findings are consistent with the contribution of astrocytes to the removal of neuronal corpses during postnatal development (Damisah et al., 2020). Lysosomal markers were also found to be increased in astrocytes of microglia-deficient *csf1r* mutant zebrafish (Berdowski et al., 2022). In summary, the gene expression evidence indicates that endosome/lysosome function in the brain is regulated to meet increased demand when microglia are absent.

### The complex impact of genetic background on viability and development of HC

**Figure S6** summarises the breeding and phenotypes of the *Fireko* on different genetic backgrounds examined in this study. C57BL/6J is the most widely used inbred mouse strain. We anticipated that the *Fireko,* having normal postnatal growth, would not replicate the lethality of the *Csf1r* knockout on this background. This was not the case. The genotype frequencies of *Csf1r*^+/+^, *Csf1r*^+/ΔFIRE^ and *Csf1r*^ΔFIRE/ΔFIRE^ mice at weaning (26%, 55%, 19%; n = 501) reported in the *Fireko*^Edin^ line (Rojo et al., 2019) are not significantly different from *Fireko*^UQ^ but the surviving *Fireko*^Edin^ mice do not develop HC. Another group recently reported the regeneration of the *Fireko* on a C57BL/6J background and found similar perinatal lethality and HC incidence to our backcrossed *Fireko*^UQ^ line (Du et al., 2026). This report raises the possibility that the improved survival in *Fireko*^UQ^ compared to *Fireko*^Edin^ backcrossed to C57Bl/6J is due either to environment (e.g. microbiome, (Caruso et al., 2019)) or breeding away from the C57BL/6JCrl substrain. The survival rate of *Fireko*^UQ^ mice and the C57BL/6J line generated by Du *et al*. (Du et al., 2026) is significantly greater than the 0.87% reported for *Csf1r^-/-^* C57BL6/J mice (Percin et al., 2018) but there is still substantial loss of *Fireko* pups due to perinatal lethality and HC. Brain ventricular enlargement is also a feature of homozygous recessive *Csf1r* mutations in FVB/NJ mice (Erblich et al., 2011) and in rats and humans (Dulski et al., 2023; Hume et al., 2020; Papapetropoulos et al., 2021). C57BL/6J mice are intrinsically susceptible to HC, and many other mutations cause HC specifically on this genetic background (Gografe et al., 2003; Goto et al., 2008; Matsumoto et al., 2020; Tapanes-Castillo et al., 2010; Wyss et al., 2012). Here we demonstrated that HC in *Fireko*^UQ^ mice is a non-communicating form. A similar probabilistic pattern of obstruction leading to lateral ventricle enlargement was reported in *Tal2*^-/-^ (Bucher et al., 2000) and *Apaf1*^-/-^ (Matsumoto et al., 2020) mice specifically on a C57BL/6J background. The mechanisms linking *Csf1r* mutations and postnatal microglial/macrophage deficiency to HC are not clear. Mutations impacting genes required for ependymal motile cilia, including *Ccdc39, Ccdc88c, Cfap221, Cfap54* and *Spef2,* lead to reduced cerebrospinal fluid (CSF) flow and postnatal HC that is also C57BL/6J strain dependent (Abdelhamed et al., 2018; McKenzie et al., 2018; Takagishi et al., 2017). Reduced numbers of parenchymal perivascular, epiplexus and intraventricular macrophages we have confirmed in neonatal *Fireko* mice might also impact CSF flow (Drieu et al., 2022; Mildenberger et al., 2022). Importantly, the acute early postnatal HC seen in *Fireko*^UQ^ mice is distinct from progressive ventricular enlargement associated, for example, with *L1CAM* mutation on the C57BL/6J background. A modifier locus on mouse Chr 5 was linked to increased penetrance of *L1CAM* mutation in C57BL/J mice (Tapanes-Castillo et al., 2010) but in *Fireko*^Edin^ mice, which do not develop HC, chromosome 5 is entirely derived from C57Bl/6J (**Figure 2E**).

The perinatal lethality and HC in C57BL/6J mice homozygous for a *Csf1r* kinase-dead mutation is also completely overcome on a C57BL/6J x BALB/c F2 background (Jacquelin et al., 2026). The impact of genetic background on other functions of microglia warrants further investigation. For example, apparently conflicting reports on the impact of the *Fireko* on synaptic pruning (O’Keeffe et al., 2024; Ramirez et al., 2026) may be a reflection of differences in genetic background between *Fireko*^Edin^ and a limited backcross to C57BL/6J (Chadarevian et al., 2024; Kiani Shabestari et al., 2022) that distinguish the two reports. Similarly, we did not replicate the changes in myelination reported in *Fireko*^Edin^ mice (McNamara et al., 2023) but myelination is also impacted by the C57BL/6J genetic background (Goudriaan et al., 2020).

### Microglial deficiency and neuropathology

The severe cognitive and motor functional deficits seen in patients with dominant mutations (Dulski et al., 2023; Konno et al., 2018; Papapetropoulos et al., 2021) are not replicated in *Fireko* mice or in mice with heterozygous disease-associated mutation. This may reflect a requirement for an environmental trigger or genetic epistasis in the human disease (Hume, 2025; Hume and Irvine, 2026). Calcification, a cardinal feature of human CRL (Dulski et al., 2023; Konno et al., 2018; Papapetropoulos et al., 2021) represents an exacerbation of a process that occurs normally in aged mice (Maheshwari et al., 2022; Wang et al., 2023). In the 5xFAD model of Alzheimer’s disease, vascular calcification in *Fireko* mice was associated with dysregulation of the *Pdgf/Pdgfrb* axis (Kiani Shabestari et al., 2022). In the thalamus of *Fireko* mice on both inbred and mixed background, astrocytosis preceded detectable calcification. Microglial depletion or *Trem2* deletion was reported to exacerbate vascular calcification in a model of primary familiar brain calcification (Zarb et al., 2021). Many of the genes implicated in PFBC are expressed by astrocytes (Maheshwari et al., 2022; Maheshwari et al., 2023; Zarb et al., 2019) but none were dysregulated in the *Fireko* thalamus. Hence, the relationship between microglial deficiency and calcification most likely reflects the loss of the essential clearance function mediated by microglial TREM2. Restoration of *Trem2* expression therefore provides sufficient explanation for the effect of BMT on thalamic calcification.

### Repopulation of the brain with bone marrow-derived microglia-like cells

Intraperitoneal transfer of WT bone marrow cells into *Fireko*^UQ^ mice at weaning leads to population of the brain with microglia-like cells (Liu et al., 2026). Flow cytometry analysis revealed that donor-derived cells were uniformly CD11b^low^/CD45^high^ resembling BAM rather than CD11b^high^/CD45^low^ microglia (Liu et al., 2026). *Adgre1* mRNA is expressed at 10-fold higher levels in BAM compared to microglia (Van Hove et al., 2019) and F4/80 was similarly elevated in BM-derived microglia by flow cytometry (Liu et al., 2026) and by IF in the current study. The functional significance of this observation is not clear. ADGRE1 structure and expression are extremely divergent across mammalian species (Waddell et al., 2018) and *Adgre1* mRNA was not selectively expressed by bone marrow-derived microglia-like cells in the rat (Carter-Cusack et al., 2024)

Repopulation of resident macrophages following BMT is not associated with detectable donor-derived monocytes in the bone marrow, blood or spleen (Liu et al., 2026). This is not entirely surprising since the *Fireko* mice are not monocyte-deficient (Liu et al., 2026) and recipients received no myeloablative conditioning to enable engraftment of bone marrow progenitors. In peripheral locations such as the peritoneum and kidney, tissue-resident macrophages were donor-derived whereas monocyte-derived macrophages remained of recipient origin for longer than 6 months post BMT (this study and (Liu et al., 2026)). At intermediate stages of repopulation of the brain, large foci of donor cells were detected apparently migrating in a wave front (Liu et al., 2026). These observations and related studies in the rat *Csf1rko* (Carter-Cusack et al., 2024) lead us to the view that the engrafting cells are progenitors with high proliferative potential that are generated in the peritoneal cavity and effectively metastasise to the brain and other organs. We speculate that engraftment may involve the clonogenic Ly6C^high^ committed monocyte progenitors identified in bone marrow which could populate tissue macrophages following intravenous adoptive transfer into macrophage-depleted recipients (Hettinger et al., 2013). The ability of the peritoneal cavity to sustain ectopic hematopoiesis has been recognized previously (Muench et al., 2011). The macrophage-deficient cavity of the *Fireko* mouse may provide a unique niche favoring macrophage progenitors. Donor cell engraftment in the cavity was not observed when BM cells were injected IP in WT mice (Liu et al., 2026) indicating that a vacant niche is essential.

There is extensive interest in microglial replacement as a treatment for neurodegenerative diseases (Rao et al., 2025) and some concern that the altered transcriptome of BM-derived microglia might promote pathology (Du et al., 2026). Published transcriptomic analyses of engrafted microglia-like cells derived from monocytes (Bastos et al., 2025; Bennett et al., 2018; Cronk et al., 2018; Du et al., 2026; Shemer et al., 2018) have used bulk or scRNA-seq analysis of isolated microglia to dissect differences between host and graft-derived cells. The universal observation in these studies is that donor-derived cells do not fully recapitulate the phenotype of resident microglia. In particular, they do not express the transcriptional repressor, *Sall1.* Although *Sall1* is enriched in microglia, it is also expressed by neurons and other glia (Munro et al., 2024). In our bulk RNA-seq data, the loss of microglia in *Fireko* mice was associated with 30-40% reduction in *Sall1,* which was not restored by BMT (Figure 6H).

The set of transcripts that was expressed selectively in the brain of BMT recipients is largely consistent with the published reports of the transcriptomic profile of bone marrow-derived microglia, supporting the validity and sensitivity of bulk RNA-seq as an alternative to tissue disaggregation that avoids the potential artefacts of scRNAseq (Hume et al., 2023). Du *et al*. (Du et al., 2026) compared the transcriptomic profile of monocyte-derived microglia in their model with previously published data. We extended that comparison to include the set of transcripts differentially detected in *Fireko*+BMT compared to WT thalamus and cortex. **Table S6** contains a Venn diagram comparing this set with DEG identified by Du *et al*. (Du et al., 2026). The overlapping set of 500 transcripts includes many of the markers discussed above. Repopulation of the brain by monocyte-derived cells was dependent upon CCR2 (Du et al., 2026), whereas rescue of the *Csf1rko* by IP adoptive transfer of bone marrow cells in neonates was CCR2-independent (Bennett et al., 2018). By contrast to its detection in monocyte-dependent models, *Ccr2* mRNA was undetectable in the *Fireko-*BMT brains.

Instead, we detected greatly increased *Cxcr4,* which is expressed by HSC-derived microglia in disease models (Werner et al., 2020) as well as the HSC marker *Hoxb4* (Antonchuk et al., 2002). Expression of the growth factor, *Il34,* was increased in response to microglial depletion with the kinase inhibitor PLX5622 and IL34 was required for repopulation by monocytes (Du et al., 2026). By contrast, neither *Il34* nor *Csf1* mRNA was altered in any brain region in the *Fireko,* although both proteins would accumulate due to the absence of microglia. PLX5622 may have off-target effects in both the brain and liver (Cao et al., 2026; Profaci et al., 2024). Similarly, we did not detect increased expression of pro-inflammatory markers, *Ifit3, Cd83* or *Il1b*, reported in monocyte-derived microglia (Du et al., 2026). Transcripts enriched specifically in the *Fireko*+BMT, including *Itgax* (CD11c), *Lgals3, Gpnmb*, *Plau* and *Igf1*, are shared with microglia engaged in prenatal and early postnatal morphogenic processes in mice ((Lawrence et al., 2024) and references therein). *Mrc1*, encoding CD206, was highly expressed in *Fireko*+BMT cortex and thalamus compared to WT but CD206 was detected in both WT and *Fireko*+BMT brains only in IBA1^-^ meningeal cells and in astrocytes (Burudi et al., 1999). The suggestion that CD206 is expressed transiently by recruited cells (Du et al., 2026) is difficult to reconcile with mRNA detection. One alternative explanation is post-translational cleavage of the receptor. The extracellular domain of MRC1 is subject to ectodomain cleavage in response to stimulation of CLEC7A (Gazi et al., 2011) which is highly expressed by the recruited cells.

We have previously reported the impact of *Csf1r* null mutation and microglial replacement following IP BMT without myeloablative conditioning in the rat (Carter-Cusack et al., 2024). As in the mouse, donor-derived microglia-like cells displayed a distinct gene expression profile compared to homeostatic microglia but we found no evidence of neuropathology and expression of stress-related genes in multiple brain regions at the time of BMT was reversed Two recent preprints (Blurton-Jones et al., 2025; Wang et al., 2025) have described neuropathology in a humanized *Fireko* mouse model following engraftment of human monocytes, including demyelination and selective loss of cortical neurons. In both cases, the distinct damage-associated microglial signature, notably an interferon signature, was much more extensive than observed in the mouse models. None of these pathologies were evident in our mice or in similar experiments by other groups (Bastos et al., 2025; Du et al., 2026).

In summary, detailed analysis of microglia-deficient mice extends the previous evidence (Hume, 2025) that microglial functions in postnatal development of the brain are largely redundant and indicates compensatory regulation of endocytic and trophic function in other cell types. Microglial deficiency in *Fireko* mice leads to age-dependent disease-relevant pathology consistent with a primary neuroprotective function and a constraint on astrocyte activation. Notwithstanding the difference in transcriptomic profile between BM-derived and resident microglia, the current study and previous analysis of BMT via IP adoptive transfer in *Csf1rko* mice and rats (Bennett et al., 2018; Carter-Cusack et al., 2024) argue that recruited cells are as effective as transplanted microglia (Chadarevian et al., 2024; Munro et al., 2024) in mitigating the pathological impacts of microglial deficiency. Our findings raise the possibility that adoptive transfer of bone marrow cells into a macrophage-depleted cavity could provide a safe and effective alternative to direct intracerebral injection or myeloablative transplantation approaches to microglial replacement (Chadarevian et al., 2024; Wu et al., 2025) especially in the context of CSF1R-related leukoencephalopathy.

## Materials and Methods

### Animals

All animal experiments in Australia were approved by The University of Queensland Health and Sciences Ethics Committee and performed in accordance with the Australian Code of Practice for the Care and Use of Animals for Scientific Purposes and Queensland Animal Care and Protection Act (2001). All experiments in Edinburgh were performed under licenses approved by the UK Home Office according to the Animals (Scientific Procedures) Act having been approved by the University of Edinburgh Local Ethical Review Board. Animals were housed in individually ventilated cages with a 12 h light/dark cycle, and food and water available ad libitum.

The generation and characterisation of *Csf1r*^ΔFIRE/ΔFIRE^ mice on a mixed C57BL/6J x CBA/J background (3:1) was described previously (Rojo et al., 2019). Gene expression profiling was performed in Edinburgh on the original genetic background. Prior to import into Australia, the *Csf1r*^ΔFIRE^ allele was backcrossed 10x to C57BL/6JCrl. To enable visualisation of myeloid populations in tissues, the imported *Csf1r*^ΔFIRE^ line was rederived and cross ed with C57BL/6JArc, then bred to the *Csf1r*-EGFP reporter transgenic line (Sasmono et al., 2003) also backcrossed >10 times to the C57BL/6JArc genetic background. For the generation of F2 intercross lines, CBA/J and BALB/cJ mice were obtained from the Animal Resource Centre, Western Australia. The *Csf1r*-FusionRed reporter transgenic line generated on the C57BL/6J background was described previously (Grabert et al., 2020). *Cx3cr1-*EGFP mice (Jung et al., 2000) were provided by Dr. Liviu Bodea (University of Queensland). Specifics of mice group size and age for each experiment are provided in respective figure legends.

### DNA Sequencing and Genotyping

DNA was extracted from tail or ear clips using the Qiagen DNeasy Blood and Tissue kit. DNA was supplied to Australian Genome Research Facility (AGRF) sequencing facility, Melbourne, Australia. Median autosomal coverage ranged from 31X to 54X. The position of all non-reference reads (where total reads was ≥ 10) was plotted against read depth. This output shows the density of non-variant reads across the genome. Genotyping of 2 male and 2 female *Fireko*^Edin^ mice was performed using the MiniMUGA genetic test by Transnetyx.

### RNA Purification

Bulk tissue RNA was isolated from two-week-old and six-week-old *Fireko*^Edin^ and WT control mice as described previously (Rojo et al., 2019). Briefly, brains from saline perfused mice were dissected, snap frozen, and disrupted in the Precellys24 Homogenizer® (Bertin Instruments). RNA isolation was performed using the RNeasy Plus Mini kit (QIAGEN).

For RNA isolation from 30-week-old mice, thalamus and cortex were dissected, snap frozen and disrupted in a FastPrep 5G Homogenizer (MP Biomedicals). RNA was isolated using TRIzol Reagent according to the manufacturer’s instructions. RNA quantity was measured on a Nanodrop Microvolume Spectrophotometer (ThermoFisher), and RNA integrity was measured on an Agilent 2100 Bioanalyzer.

### Library preparation and sequencing

For samples from juvenile mice, library preparation and paired-end RNA sequencing was performed by Edinburgh Genomics. The average read depth per sample was 59.0 million reads (median 57.3 M), with a minimum of 47.0 M and a maximum of 91.5 M. RNA-sequencing reads were mapped to the mouse reference genome using the STAR RNA-seq aligner. Tables of per-gene read counts were generated using featureCounts. Further details are available at (https://www.ebi.ac.uk/biostudies/arrayexpress/studies/E-MTAB-14156).

Library preparation for 30-week-old mice was performed at the University of Queensland Sequencing Facility (University of Queensland, Brisbane, Australia) and the sequencing was done in conjunction with services through the Australian Genome Research Facility (Australia). Briefly, RNA-Seq libraries were generated using the Illumina Stranded mRNA Library Ligation kit (Illumina, 20040534) and Illumina RNA UD Indexes Ligation (Illumina, 20091657) according to the standard manufacturer’s protocol (Illumina, Document # 1000000124518 v04, Oct 2023). 80 ng of total RNA was enriched for mRNA using an oligo-dT bead isolation method, and the mRNA was fragmented in a heat fragmentation step.

cDNA was synthesized from the fragmented RNA using random primers. The first strand cDNA was converted into dsDNA in the presence of dUTP to prevent subsequent amplification of the second strand and thus maintaining the strand orientation of the original mRNA. The 3’ ends of the cDNA were adenylated and pre-index anchors were ligated. The libraries were amplified with 13 cycles of PCR, incorporating unique indexes for each sample to produce libraries ready for sequencing. The libraries were quantified on the Revvity LabChip GX Touch with the DNA High Sensitivity Reagent kit (Revvity, CLS760672). Libraries were pooled in equimolar ratios.

Sequencing was performed using the Illumina NovaSeq X Series sequencer (NovaSeq Control Software v1.3.159007 and Real Time Analysis v4.29.3). The library pool was diluted and denatured according to the standard NovaSeq X protocol and sequenced to generate paired-end 151 bp reads using a NovaSeq X Series 10B Reagent Kit (300 cycles) (Illumina, 20085594). After sequencing, fastq files were generated using bcl2fastq2 (v2.20.0.422), and sequencing adapters were trimmed. Raw data and further details are available at (https://www.ebi.ac.uk/biostudies/arrayexpress/studies/E-MTAB-17673).

The nf-core/rnaseq bioinformatics pipeline was used to analyse sequencing data from 30-week-old animals, using the GRCm39 Mus musculus reference transcriptome. Transcript expression level files were then imported into Visual Studio Code (v1.136.1) where all downstream analysis was conducted using R. Differential expression analysis was conducted with DESeq2, using genotype and BMT treatment as a combined design model. Genes where there were less than three samples with normalized counts greater than 10 were filtered out.

Principal component plots were created using the plotMDS function in R. Volcano plots were created using the ggplot function in R, and the lfcShrink function from DESeq2 was used with the ‘apeglm’ method to shrink log_2_ fold changes to reduce the effects of outliers. Bar graphs were created in GraphPad Prism (10.3.0) from averaged read counts.

### Network cluster analysis of gene expression

BioLayout (http://biolayout.org) and Graphia (https://graphia.app) each calculate a Pearson correlation matrix comparing similarities in gene expression profiles between samples to generate a network graph. For the more complex visualization of multiple brain regions from juvenile mice, we used Biolayout, whereas visualization of the BMT data in Figure 6 used Graphia. Only relationships where r > 0.85 were included. A network graph was created in which nodes (genes) were connected by edges (representing correlation coefficients) using the Markov Cluster Algorithm (MCL) with an inflation value of 2.0.

To generate the heatmap in Figure 1, raw gene expression data was averaged by genotype (WT n=3, KO n=4) for each region. Each gene was then z-score normalised across samples and values capped at ± 2. Data were visualised using a heatmap with the pheatmap package (v1.0.12) in R (v4.4.1).

### Tissue disaggregation and flow cytometry

Brain tissues were processed using enzymatic digestion and density gradient centrifugation. Collagenase Type IV (10 mg) was prepared in 10 mL HBSS, supplemented with 20 µL DNase I (10 mg/mL) and 1 mg dispase per 10 mg collagenase. Mice were euthanized, and brains were dissected, halved sagitally, and minced. The tissue was incubated in collagenase on a rocking platform at 37°C for 45 min, before being filtered through a 70 µm strainer. After centrifugation at 500 x g for 5 min at 4°C, the pellet was resuspended in 12.5 mL isotonic Percoll and centrifuged at 800 x g for 45 min at 4°C (no brake). The myelin layer and supernatant were removed, and the pellet was washed twice with FACS buffer. The final pellet was resuspended in FACS buffer.

Peripheral blood was collected via retro-orbital bleed into EDTA-coated tubes and analysed using an automated hematology analyser, Mindray BC5000 Haematology Analyser (TRI Flow Cytometry Facility). Blood was then subjected to red blood cell lysis for 10 min in lysis buffer (150 mM NH_4_Cl, 10 mM KHCO_3_, 0.1 mM EDTA, pH 7.4), followed by two PBS washes and resuspension in FACS buffer for staining. Peritoneal cells were collected by lavage with 10 mL of PBS and centrifuged at 400 x g for 5 min at 4°C before incubated in red blood cell lysis buffer for 10 min. Cells were then pelleted again before resuspension in 1 mL of FACS buffer for staining. Bone marrow (BM) cells were prepared by flushing femurs and tibias with 10 mL of PBS, then centrifuging at 400 x g for 5 min at 4°C. The pellet was incubated in red blood cell lysis buffer for 10 min, before resuspension in 1 mL of FACS buffer for staining.

Once a single cell suspension had been obtained, cells were stained with fluorophore-conjugated antibodies on ice and analysed on an LSRFortessa (BD). Data were analysed using FlowJo v10 Software (BD Life Sciences, NJ, USA).

### Flow Cytometry antibodies

| **Antibody** | **Fluorochrome** | **Supplier** | **Catalogue Number** |
| --- | --- | --- | --- |
| Anti-mouse/human CD11b | PerCP-Cy5.5 | Biolegend | 101228 |
| Anti-mouse/human CD11b | BV510 | Biolegend | 101263 |
| Anti-mouse Ly-6G | BV785 | Biolegend | 127645 |
| Anti-mouse Ly-6C | Pacific Blue | Biolegend | 128014 |
| Anti-mouse Ly-6C | PerCP-Cy5.5 | Biolegend | 128012 |
| Anti-mouse CD117 (c-kit) | PECy7 | Biolegend | 135112 |
| Anti-mouse CD115 | BV605 | Biolegend | 135517 |
| Anti-mouse F4/80 | BUV737 | BD | 749283 |
| Anti-mouse F4/80 | AF647 | Biolegend | 123122 |
| Anti-mouse CD45 | BUV395 | BD | 564279 |
| Anti-mouse Ly-6A/E (Sca-1) | BV650 | Biolegend | 108143 |
| Anti-mouse CD135 | APC | Biolegend | 135310 |
| Anti-mouse CD48 | Pacific Blue | Biolegend | 103418 |
| Anti-mouse CD150 | BV711 | Biolegend | 115941 |
| Anti-mouse NK1.1 | AF700 | BD | 560515 |
| Biotin anti-mouse CD5 | NA | Biolegend | 100604 |
| Biotin anti-mouse CD3 | NA | Biolegend | 100244 |
| Biotin anti-mouse TER-119 | NA | Biolegend | 116204 |
| Biotin anti-mouse/human<br>CD45R/B220 | NA | Biolegend | 103204 |
| Streptavidin | APC-Cy7 | Biolegend | 405208 |

### Immunohistochemistry, immunofluorescence, and imaging

For free-floating sections, brains were dissected from the skull and immersion fixed in 4% PFA for 5 hours at room temperature (RT), washed once in PBS and stored in PBS with 0.1% Sodium Azide. Brains were serially sectioned at 30 μm in the sagittal or coronal plane using a Leica VT1200S vibratome, and sections were transferred to a 24-well-plate and washed in PBS before staining. Sections were incubated in permeabilization buffer (1% Triton-X, 0.1% Tween-20 in PBS) for 2 h at RT, followed by blocking buffer (4% serum, 0.3% Triton-X, 0.05% Tween-20 in PBS) for 2 h at RT then in primary antibody diluted in blocking buffer overnight at 4◦C on a rocking platform. Primary antibodies included rabbit anti-IBA1 (1:500, Novachem, 019-19741), chicken anti-IBA1 (1:500, Synaptic Systems, 234 009), rabbit anti-GFAP (1:500, Dako, Z0334), chicken anti-GFAP (1:500, Invitrogen, PA1-10004), guinea pig anti-TMEM119 (1:500, Synaptic Systems, 400 004), rat anti-MBP (1:500, abcam, ab7349), mouse anti-neurofilament marker (SMI312, 1:1000, Biolegend, 837904), rabbit anti-CD206 (1:500, Abcam, ab64693), rat anti-Galectin-3 (1:200, Invitrogen, 14-5301-85), guinea pig anti-parvalbumin (1:500, Synaptic Systems, 195 004). To label F4/80, antigen retrieval in Tris/EDTA solution was performed on free-floating sections for 10 min before permeabilization, blocking, and incubation in primary antibody rabbit anti-F4/80 (1:500, Abcam, ab300421). To label perineuronal nets, brain sections were co-stained with *Wisteria floribunda* agglutinin (WFA; Merck, L1516) diluted 1:500 in blocking buffer. To label thalamic calcification, brain sections were co-stained with AF647-RIS Imaging Reagent (BioVinc LLC, BV500101) diluted 1:500 in blocking buffer. After overnight incubation, brain sections were washed 3 x 5 min with PBS then incubated for at least 2 h in appropriate fluorophore-labelled secondary antibodies (Donkey anti-rabbit AF647 (1:500, Invitrogen, A31573); goat anti-chicken AF594 (1:500, Invitrogen, A32759); donkey anti-rabbit AF488 (1:500, Invitrogen, A21206); goat anti-rat AF647 (1:500, Invitrogen, A48265); goat anti-chicken AF488 (1:500, Invitrogen, A32931); goat anti-rabbit AF594 (1:500, Invitrogen, A32740); donkey anti-guinea pig CF633 (1:500, Merck, SAB4600129); goat anti-rabbit AF750 (1:500, Invitrogen, A21039); goat anti-mouse AF647 (1:500, Invitrogen, A21235)) diluted in blocking buffer. Sections were once again washed 3 x 5 min with PBS before nuclei were stained with DAPI (1:5000, Thermo Scientific, 62248) for 5 min. Sections were washed a final time and mounted onto glass slides with Fluorescence Mounting Medium (Dako). Free floating sections were also incubated in 1% w/v Alizarin Red S (1% Alizarin Red S Sigma Cat# A5533 in RO water, pH 5.5) with orbital agitation for 10 min at RT. Sections were washed with PBS 3 x 5 min and mounted using Dako fluorescence mounting media.

For paraffin-embedded sections, brains were sectioned at 6 µm using a Leica RM2245 microtome. After dewaxing and rehydrating the tissues, endogenous peroxidases were quenched by immersion in 4% H_2_O_2_ in methanol for 5 min. For detection of astrocytes, sections were incubated overnight with primary antibody rabbit polyclonal anti-GFAP (Dako, Glostrup, Denmark) diluted 1:2000. Primary antibody binding was detected using biotinylated goat anti species-specific antibodies (Jackson Immunoresearch, Cambridge, UK) and visualised using the Elite ABC/HRP kit (Vector Laboratories, Peterborough, UK) and diaminobenzidine (DAB) (Merck, Glasgow, UK) between stringent washing steps. Sections were lightly counterstained with hematoxylin. For the detection of endogenous calcium, deparaffinised sections were immersed in 2% Alizarin Red S solution in water (pH 4.1) for 5 min followed by quick rinses in acetone, 50:50 acetone-xylene and finally clarified in xylene.

Neonatal mice were euthanised by decapitation before their brains were dissected and fixed in 4% PFA overnight at RT. Brains were washed once in PBS before cryoprotection in 30% sucrose solution for 48 hours and frozen in OCT. Brains were sectioned coronally at 40 μm using a cryostat. Sections were fixed again in 4% PFA for 10 min, before being immersed in ice-cold methanol for 10 min. Blocking buffer was applied for 2 hours before sections were incubated in primary antibodies rabbit anti-IBA1 (1:500, Novachem, 019-19741) and chicken anti-GFAP (1:500, Invitrogen, PA1-10004) diluted in blocking buffer, overnight at 4. Sections were washed 3 x 5 min with PBST before incubation in secondary antibodies donkey anti-rabbit AF647 (1:500, Invitrogen, A31573); goat anti-chicken AF594 (1:500, Invitrogen, A32759) diluted in blocking buffer, for 2 hours at RT. Sections were washed 3 x 5 min again and DAPI stained before they were cover-slipped.

### Evans Blue intracerebroventricular injection

To investigate the nature of HC, mice were anaesthetised using isoflurane and the superior side of the skull was exposed. Mice then received intracerebroventricular injection of Evans Blue dye. Mice were kept anaesthetised for 10 min to allow the flow of cerebrospinal fluid before euthanasia, extraction of the brain and visualisation of dye within the ventricular system.

### Behaviour analysis

The novel object recognition, marble burying and elevated plus maze tests (Lueptow, 2017; Patkar et al., 2019) were each applied to the same cohort of WT, *Csf1r*^ΔFIRE/+^ and *Fireko* mice (C57BL6/J x CBA/J F2) at 30 weeks of age prior to sacrifice for tissue collection and imaging. The gait of unforced movement was analysed in *Fireko*^Edin^ mice at weekly intervals from 8 weeks of age with the CatWalk^TM^ XT system (Noldus Information Technology, Wageningen, The Netherlands). CatWalk XT was operated in a dedicated dark and quiet room and habituation and experimental procedures were performed during the same period between 9am and 3pm. Compliant runs were between 1.5 and 6 seconds.

### Mouse brain magnetic resonance imaging (MRI)

30-week-old C57BL6/J x CBA/J F2 mice were perfuse-fixed with 0.1M Phosphate Buffer with 1% EM-grade glutaraldehyde. Brains were subsequently fixed for 3 h at RT then transferred to PBS with 0.01% sodium azide until magnetic resonance imaging (MRI). Fixed brains were washed in PBS with 0.2% v/v gadopentetate dimeglumine (Magnevist, Bayer, Leverkusen, Germany) for 4 days to shorten the T_1_ relaxation and enhance MRI contrast (Kurniawan, 2018). MRI data were acquired using a 16.4 T vertical bore microimaging system (Bruker Biospin, Rheinstetten, Germany; ParaVision v6.01) equipped with Micro2.5 imaging gradient and a 15 mm linear surface acoustic wave coil (M2M, Brisbane, Australia). Three-dimensional (3D) T_1_/T_2_*-weighted gradient echo images were acquired with the following parameters: repetition time (TR) = 50 ms, echo times (TE) = 12 ms, bandwidth = 50 KHz, field of view (FOV) = 19.6 × 12 × 9 mm and matrix size = 654 × 400 × 300, which resulted in 30 μm 3D isotropic image resolution and image acquisition time of 56 mins. 3D diffusion-tensor image (DTI) data were acquired using a Stejskal-Tanner DWI spin-echo sequence with TR = 200 ms, TE = 23 ms, δ/Δ = 2.5/12 ms, bandwidth = 50 KHz, FOV =19.6 × 12 × 9 mm and matrix size = 196 × 120 × 90, image resolution = 100 μm, 30 direction diffusion encoding with b-value = 5000 s/mm^2^, two b = 0 images, with the acquisition time of 16 h 2 min.

Susceptibility-weighted imaging (SWI) was generated for the mouse brain data using Paravision 6.0.1. The anatomical images were constructed by combining multi-echo images to enhance image contrast. Images were registered into the Waxholm Space MRI/DTI template and the C57BL/6J mouse brain MRI atlas (Ma et al., 2005) using ANTs diffeomorphic image registration (Avants et al., 2011). The model-based segmentation of brain regions was performed on each sample and their image volumes and intensities were measured using ITK-SNAP.

### MRI data analysis

The FID of DTI datasets were zero-filled by a factor of 1.5 in all dimensions prior to Fourier transform to improve fibre tracking. DTI data were bias corrected using ANTs N4BiasFieldCorrection and processed using MRtrix3 software (www.mrtrix.org). Fiber orientation distribution (FOD) was reconstructed using constrained spherical deconvolution (CSD) method, and probabilistic tractography was performed using iFOD2 algorithm. Tractography was performed for specific major white matter (WM) tracts. Firstly, the seeding regions of interest (ROIs) were manually drawn in the midsagittal and coronal sections of the colour vector map, and fibre tracks were generated for the corpus callosum, hippocampal commissure, internal capsule and anterior commissure at 100 seeds per voxel. The volumes of the white matter tracts were measured from Tract Density Intensity maps (Calamante et al., 2012) with the intensity threshold set at 20% to remove background noise from probabilistic tracking.

### Image quantification and analysis

For confocal microscopy, FV3000 Confocal and Nikon Crest-Optics Spinning Disc Confocal microscopes were used to acquire images using 4x – 40x objectives. Quantification of IBA1, GFAP, and TMEM119 in Figure 3 was performed using (Fiji Is Just) ImageJ (Fiji; v2.14.0). The percentage area of positive staining was calculated using the ‘Measure’ tool following adjustment of brain region-specific threshold. The number of meningeal macrophages was counted and normalised to length of meninge. The number of choroid plexus macrophages was normalised to the average area of choroid plexus across all mice. In Figure 4, quantification of MBP used a minimum of four images per brain region for each mouse, for three mice of each genotype. For quantification of GFAP and risedronate, one region was analysed per section using colour thresholding with MaxEntropy method, with the same threshold used across three mice of each genotype. For quantification of PNNs and PV neurons, two regions of somatosensory cortex were analysed per mouse, for three mice of each genotype.

For all other microscopy quantification, quantification was performed in QuPath (https://qupath.github.io; v0.6.0). Regions of interest were annotated, before a pixel classifer was created usilising images from all genotypes. Once trained, classifiers were used to calculate the area of positive staining and number of cells across regions and the area and fluorescence intensity of each positive cell.

## Data analysis

Statistical analysis of data was performed using GraphPad Prism (v10.3.0). For comparing two groups, an unpaired Student’s t-test with Welch’s correction was applied. For comparisons involving three or more groups, ordinary one-way and two-way ANOVA followed by Tukey’s multiple comparison tests was utilized. p values < 0.05 were considered significant.

## Data availability

All processed RNAseq data are provided in Supplementary Tables. Primary data for Tables S1 and S2 is available at https://www.ebi.ac.uk/biostudies/arrayexpress/studies/E-MTAB-14156. Primary data for Tables S4, S5, and S6 is available at https://www.ebi.ac.uk/biostudies/arrayexpress/studies/E-MTAB-17673.

## Acknowledgments

The Irvine, Hume and Summers laboratories gratefully acknowledge core funding from The Mater Foundation, Brisbane, Australia. The generation, breeding and analysis of the *Csf1r*^ΔFIRE/ΔFIRE^ mice in Edinburgh was funded by a grant from the Medical Research Council (MRC) UK grant <u>MR/M019969/1</u> to DAH. Additional support was provided by the Simons Initiative for the Developing Brain and by the UK Dementia Research Institute, which receives its funding from DRI Ltd, funded by the UK Medical Research Council, the Alzheimer’s Society and Alzheimer’s Research UK. The current work was supported by Australian National Health and Medical Research Council (NHMRC) Grant <u>GNT1163981</u> awarded to DAH and KMS. DAH is supported by NHMRC (Australia) Investigator Grant 2009750. We thank Dr. Liviu Bodea for the *Cx3cr1*^GFP/+^ mice and thank Professor Allison Pettit for the anti-Galectin-3 antibody. This work was supported by resources provided by The University of Queensland Research Computing Centre’s Bunya supercomputer (https://dx.doi.org/10.48610/wf6c-qy55), with funding from The University of Queensland, Brisbane, Australia. We acknowledge input and expertise from the Biological Resources facility and the Microscopy, Histology and Flow Cytometry facilities of the Translational Research Institute (TRI) and The Roslin Institute. TRI is supported by the Australian Government. NAM was supported by project (BB/S005471/1) and Institute Strategic Programme Grant funding from the Biotechnology and Biological Sciences Research Council (grant numbers BBS/E/RL/230001C & BBS/E/RL/230002B).

## Legends to Supplementary Figures

**Supplementary Figure 1.**
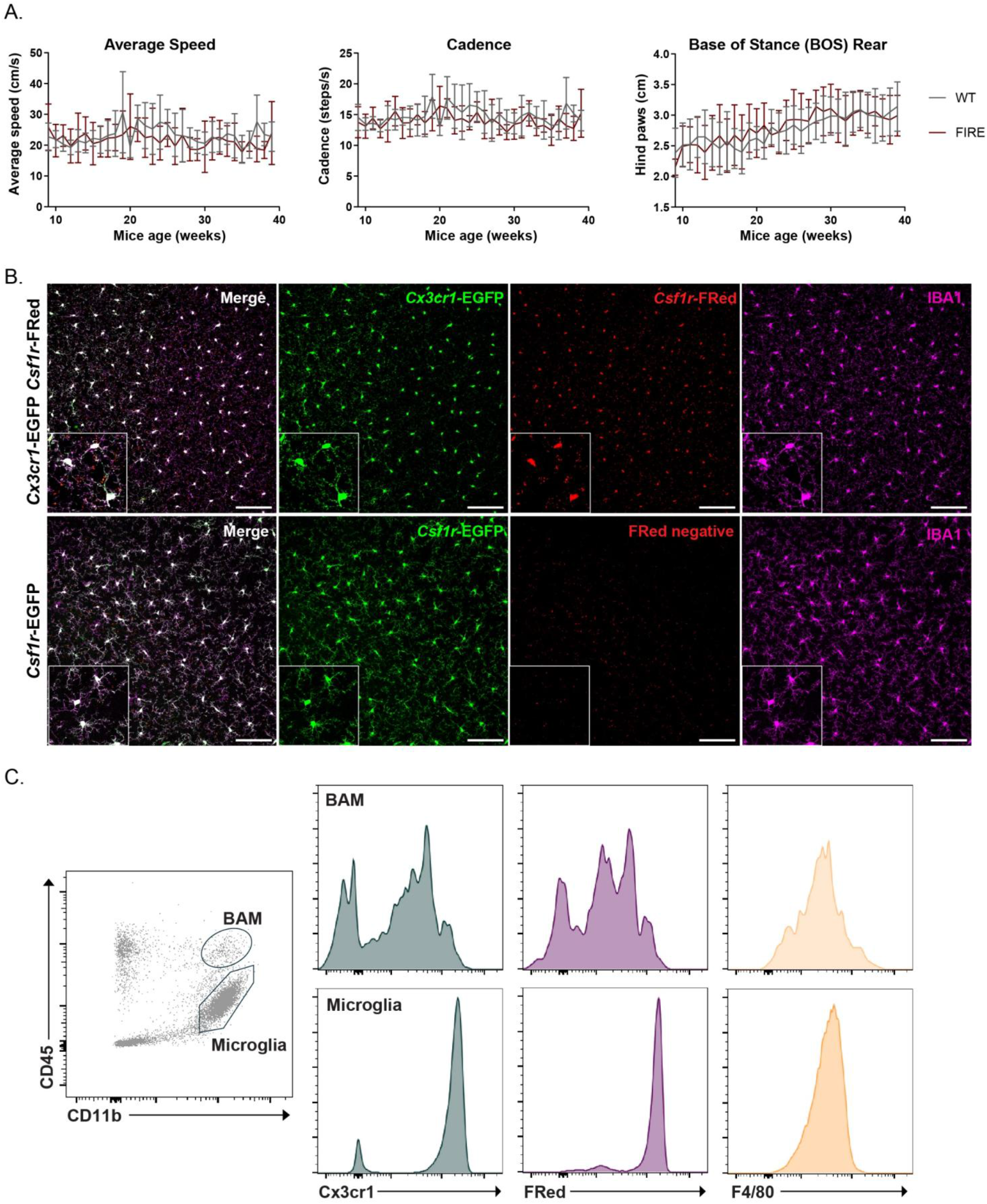
(A) Gait analysis as described in Methods was performed weekly on cages of *Fireko*^Edin^ mice and WT littermates of both sexes (n = 10) with *Csf1r* genotype blinded to the operator. For the gait parameters, three compliant runs of each mouse made up the testing run. Following acquisition of the whole data set, runs were grouped by genotype and age. No significant difference based upon genotype was detected. (B) To confirm that CSF1R expression is microglia restricted, free-floating sections of cortex from *Cx3cr1*-EGFP x *Csf1r-*FusionRed mice were stained for IBA1 and counterstained with DAPI. Note the complete overlap of the three markers. The FusionRed (FRed) reporter is detected primarily in the cell body of microglia. Lower panel shows cortex of *Csf1r-*EGFP mouse also stained for IBA1, providing a negative control for FRed. Scale bars = 100 µm. (C) Representative flow cytometry profile of total brain cells isolated from *Cx3cr1*-EGFP x *Csf1r-*FusionRed mice. Note the variable detection of *Cx3cr1*-EGFP and FRed in cells in the brain-associated macrophage (BAM) gate, likely reflecting the presence of monocytes.

**Supplementary Figure 2.**
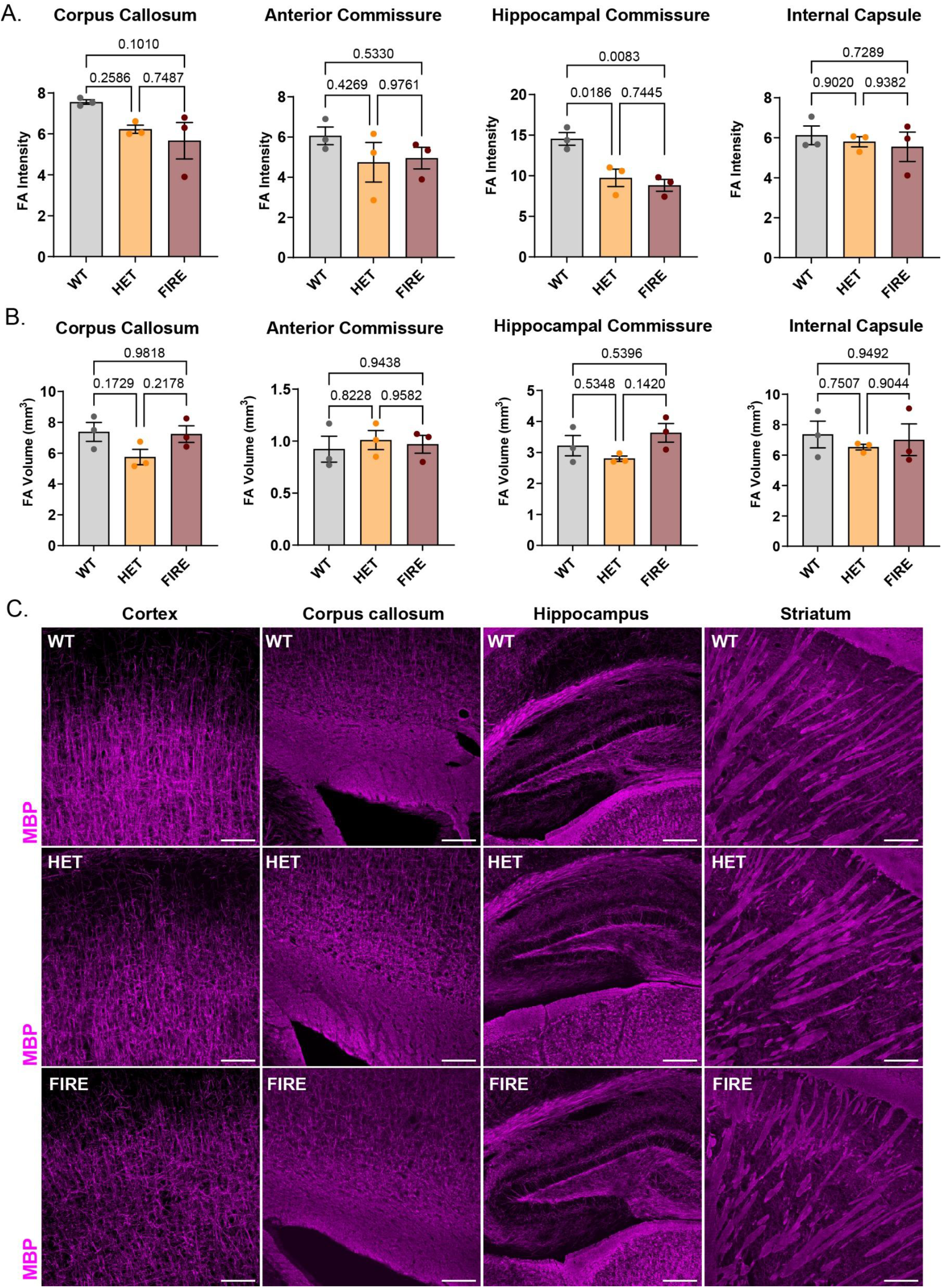
(A-B) Quantitative analysis of MRI imaging. Fractional anisotropy (FA) provides a quantitative biomarker of white matter integrity. Graphs show mean ± SEM from three mice of each genotype (see images in Figure 4D), analysed by ordinary one-way ANOVA followed by Tukey’s multiple comparisons test. P values are shown on each graph. (C) Brains of the 30-week-old *Fireko*^BCBF2^ mice and littermate WT controls were stained to detect myelin basic protein (MBP; quantification in Figure 4F). Representative images of cortex, corpus callosum, hippocampus, and striatum are shown, from three mice per genotype. Scale bars = 100 µm.

**Supplementary Figure 3.**
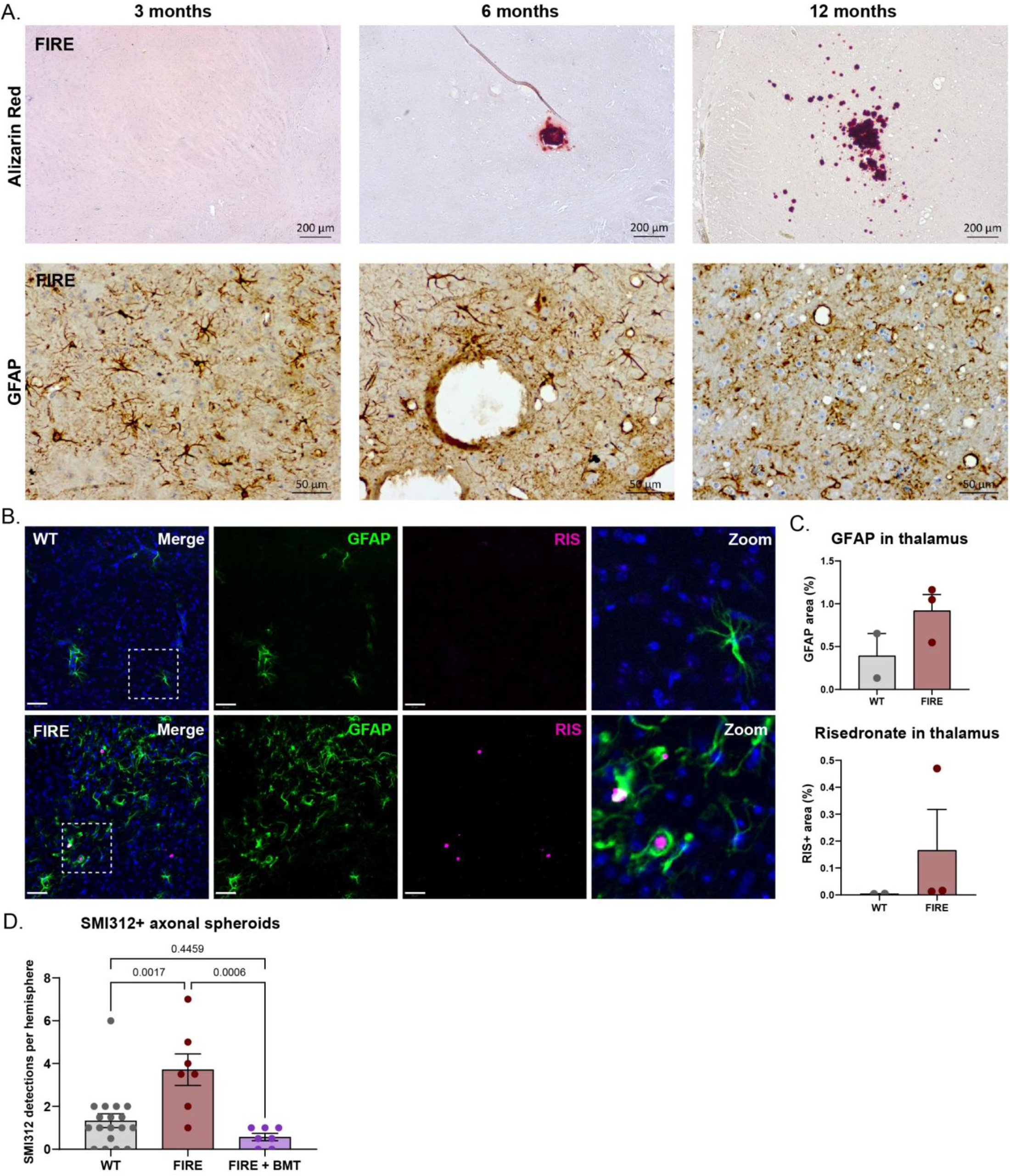
(A) Sections of thalamus from *Fireko*^Edin^ mice were stained for GFAP and calcification (Alizarin Red). In the cohort of 15 mice at 3 months, 10 showed substantial astrocytosis detected with anti-GFAP without evident calcification. Scale bars = 200 µm (top) or 50 µm (bottom). (B-C) Free-floating sections of thalamus from 15-week-old WT and *Fireko*^UQ^ mice were stained to detect GFAP and calcium deposits (risedronate). Increased GFAP was detected in all *Fireko*^UQ^ mice (n = 3). Image shows the individual *Fireko*^UQ^ mouse in which small foci of calcification were detected, intimately associated with GFAP+ astrocytes. Scale bars = 50 µm. (D) Sections of brains from *Fireko*^UQ^ mice were stained with neurofilament marker SMI312 for axonal spheroids. The number of SMI312+ spheroids per hemisphere was quantified from two sections for each animal. Graph shows mean ± SEM of at least seven mice per genotype, analysed by ordinary one-way ANOVA followed by Tukey’s multiple comparisons test. P values are shown on the graph.

**Supplementary Figure 4.**
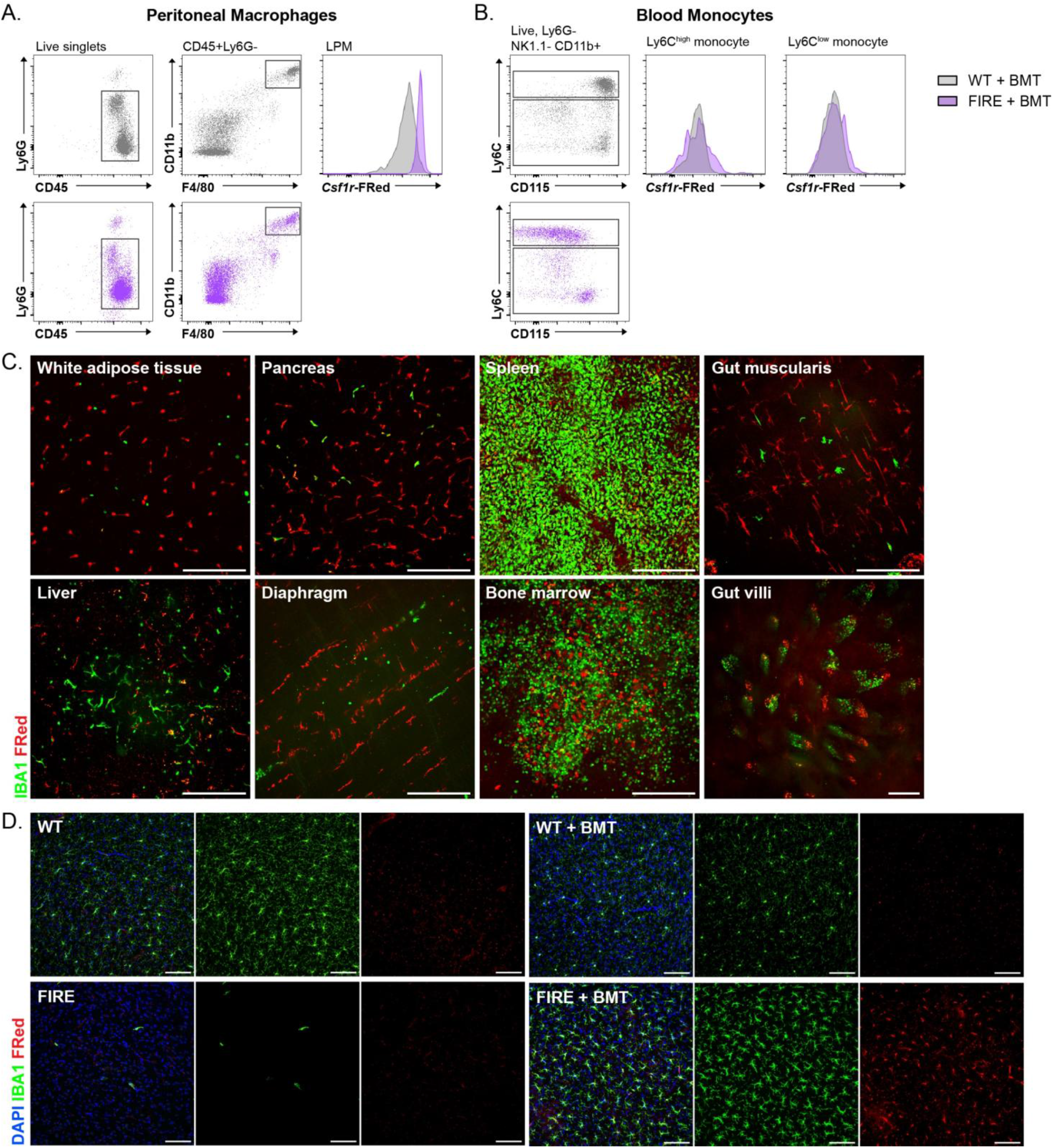
WT and *Fireko*^UQ^ mice received WT bone marrow transplant (BMT) from *Csf1r*-FusionRed (FRed) mice at 3 weeks. Untreated WT and *Fireko*^UQ^ mice act as controls. (A-B) Flow cytometry gating strategy for peritoneal macrophages (A) or blood monocytes (B), and histograms illustrating *Csf1r*-FRed expression. (C) Whole mount imaging of various tissues from *Fireko*^UQ^ mice treated with BMT. Native recipient cells were positive for *Csf1r-*EGFP, while donor-derived cells were positive for *Csf1r-*FRed. Three animals were imaged. Scale bars = 200 µm. (D) Free-floating sections of thalamus from WT and *Fireko*^UQ^ treated/untreated with BMT were stained to detect IBA1. Donor-derived cells were positive for *Csf1r-*FRed transgene. Representative images are shown (n = 5-6 from each genotype). Scale bars = 100 µm.

**Supplementary Figure 5.**
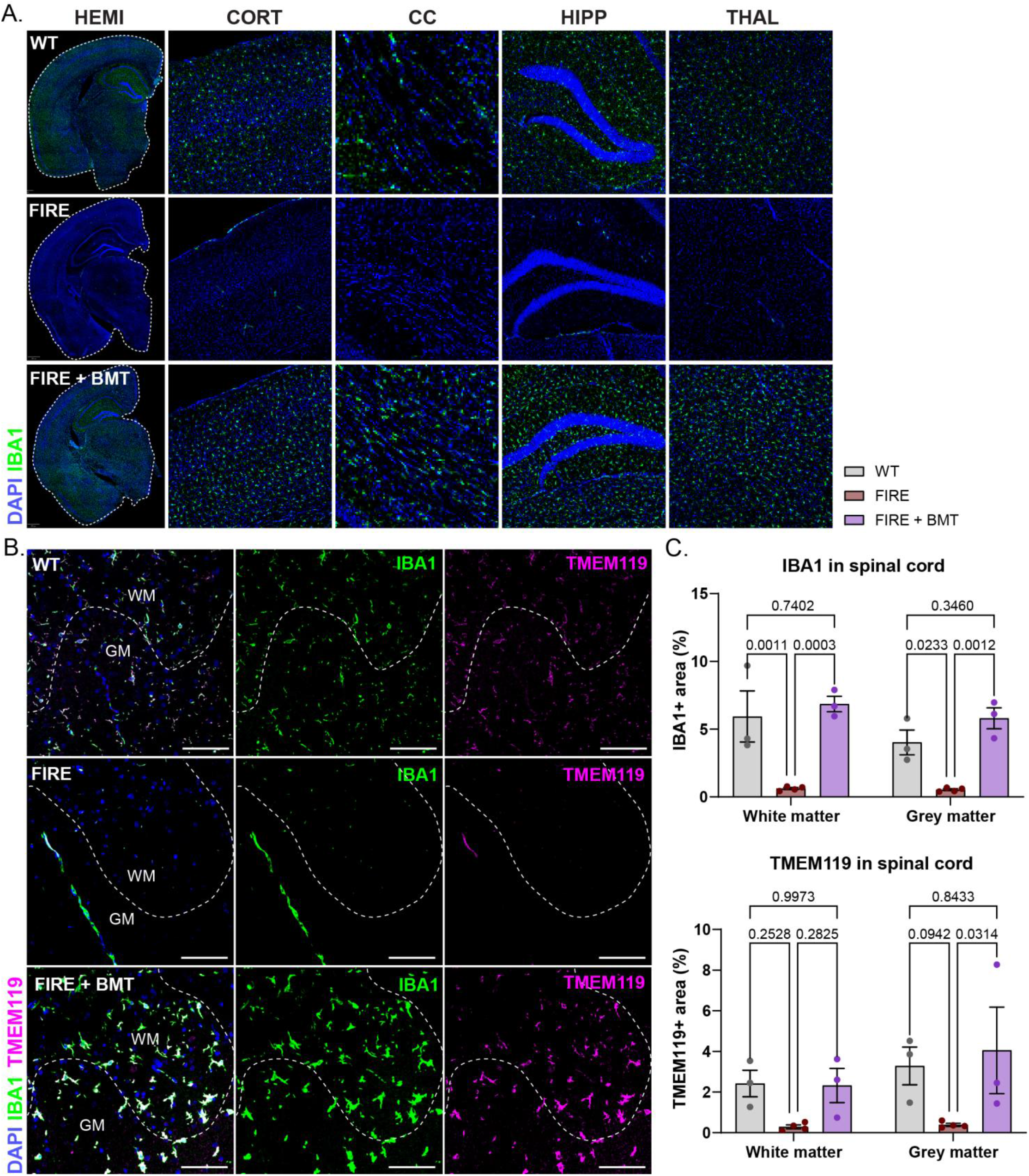
(A) Sections of brain from 30-week-old WT, *Fireko*^UQ^ (FIRE) and *Fireko*^UQ^ + BMT (FIRE + BMT) mice were stained with IBA1. Representative images of the entire hemisphere (HEMI), cortex (CORT), corpus callosum (CC), hippocampus (HIPP), and thalamus (THAL) are shown. (B) Cryosections of thoracic spinal cord from 30-week-old cohort were stained with IBA1 and TMEM119. White matter (WM) and grey matter (GM) are highlighted on images. Scale bars = 100 µm. (C) Percentage area of IBA1^+^ (top) and TMEM119^+^ (bottom) staining was quantified from at least three mice per genotype. Graphs show mean ± SEM, analysed by ordinary two-way ANOVA followed by Tukey’s multiple comparisons test. P values are shown on each graph.

**Supplementary Figure 6.**
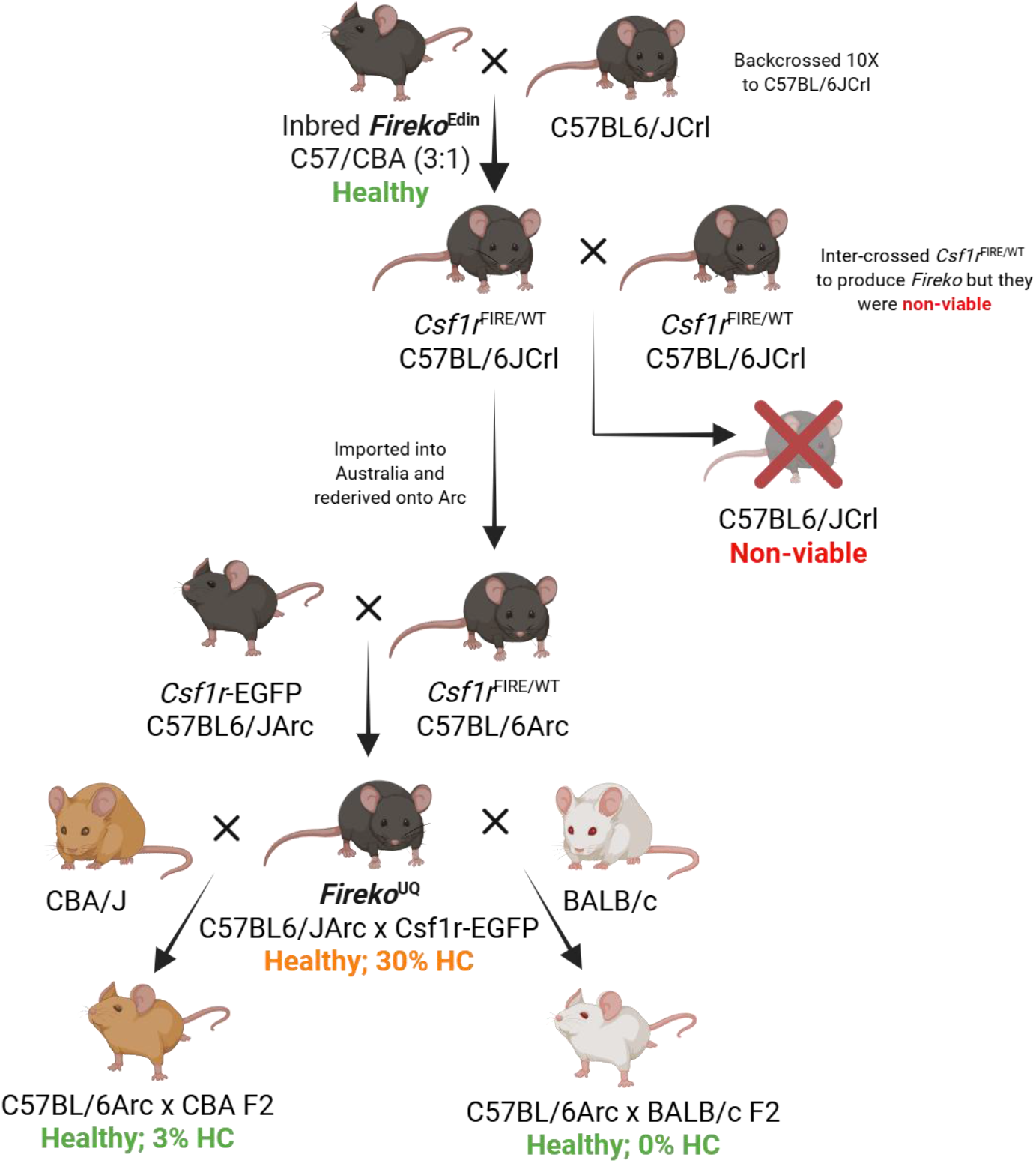
Schematic of breeding strategy undertaken to produce *Fireko*^UQ^ mice and subsequent crosses to CBA/J and BALB/c backgrounds. Viability and incidence of hydrocephalus (HC) is reported. Created in BioRender. Taylor, I. (2027) https://BioRender.com/i23mn0n

## Supplementary On-line data

**Table S1.** Total RNA-seq analysis of multiple brain regions from WT and *Fireko* mice at 2 weeks of age.

**Table S2.** Total RNA-seq analysis of multiple brain regions from WT and *Fireko* mice at 6 weeks of age.

**Table S3.** Co-expression clusters identified by network analysis of combined RNA-seq data from multiple brain regions from WT and *Fireko* mice at 6 weeks of age.

**Table S4.** Total RNA-seq analysis of thalamus from WT, *Fireko*, and *Fireko* + BMT mice at 30 weeks of age (27 weeks post BMT).

**Table S5.** Total RNA-seq analysis of cortex from WT, *Fireko*, and *Fireko* + BMT mice at 30 weeks of age (27 weeks post BMT).

**Table S6.** Differentially expressed gene (DEG) analysis comparing *Fireko* to WT, *Fireko +* BMT to WT, and Venn diagram comparing DEG list from this study to DEG list from Du *et al* (Du et al., 2026).

**Table S7.** Co-expression clusters identified by network analysis of combined RNA-seq data from thalamus and cortex from WT, *Fireko*, and *Fireko* + BMT mice at 30 weeks of age (27 weeks post BMT).

